# Analyses of protein expression and genetic fitness determinants reveal dynamic pathways active in starved *Pseudomonas aeruginosa*

**DOI:** 10.1101/2025.09.28.678986

**Authors:** Findlay D. Munro, Elize Ambulte, Claudia M. Hemsley, Megan Bergkessel

## Abstract

Heterotrophic bacteria rapidly deplete essential macronutrients during growth and must navigate subsequent periods of growth arrest imposed by starvation. Nutrient limitations can be dynamic in nature, requiring ongoing regulatory adjustments involving new protein synthesis despite total biosynthetic activities being dramatically lower than during growth. Here, we have characterized the responses of the opportunistic pathogen *Pseudomonas aeruginosa* to prolonged starvation for carbon or nitrogen sources, and to transitions between these states. We find that most cells survive both types of starvation for more than a week and maintain low but robustly detectable levels of protein synthesis in the absence of growth. Nitrogen-starved cells are larger, make more proteins and retain fewer ribosomes than carbon-starved cells, indicating that distinct physiological strategies are adopted during the two starvation types. We found that the newly synthesized proteomes of each starvation type are distinct, although many of the most highly synthesized proteins are shared between both conditions. Interestingly, we observed a temporary burst of protein synthesis as cells were transitioned between the two starvation conditions, which may reflect active remodelling of the proteome during growth arrest. We also used transposon insertion sequencing to identify genes impacting fitness in both starvation conditions and during transitions between the two and found that a highly overlapping set of global regulators most strongly influenced survival. Combining these datasets, we highlight proteases and chaperones; flagellar motility; and the nitrogen-related phosphotransferase system as key fitness-impacting functions that are actively maintained by growth arrested *Pseudomonas aeruginosa*.

**Importance:** Molecular microbiology has traditionally focused on exponential growth in model organisms as the preferred context in which to study bacterial physiology, especially the regulation of new protein synthesis. However, in natural environments, including many infection contexts, Proteobacteria frequently enter growth arrest due to nutrient limitation. The dynamics and regulation of protein synthesis in growth-arrested cells remain poorly understood, especially in pathogens. Furthermore, growth arrest increases tolerance to a variety of stresses, including many clinically used antimicrobials. We have conducted a comprehensive exploration of the proteins being made by growth arrested *Pseudomonas aeruginosa* during total nitrogen or carbon starvation and at the transition between these two starvation types, and the genes supporting fitness under these conditions. These datasets suggest dynamic redistribution of resources among important cellular functions and will serve as a resource for further investigations of starvation-induced growth arrest, a ubiquitous but understudied physiological state of heterotrophic bacteria.

## Introduction

Historically, bacterial physiology and regulation have often been studied under conditions that support balanced, steady-state, exponential growth. These efforts have permitted quantification of the relationships between growth rate, ribosome abundance, translation rate and nutrient quality (the “bacterial growth laws”, reviewed in^1,2^ and references therein) and have provided essential insights into the coordination of metabolism with gene expression during rapid growth. For example, concentrations of proteins in the *E. coli* proteome are determined almost entirely by transcription initiation rates at their promoters^3^, and the fraction of the cell’s mass that is comprised by ribosomes is linearly related to the growth rate^4^. Rapid, steady state growth also offers practical advantages for studying responses to perturbations. During rapid growth high rates of metabolic activity, gene expression and proteome turn-over can power robust and easily measured responses.

However, many observations from natural environments suggest that bacteria frequently exist in physiological states outside of steady-state growth. Bacteria that grow in association with a host are often constrained by the host, leading to growth rate estimates that are sometimes surprisingly slow or highly variable^5–7^. Population blooms of organisms that were scarce before nutrient amendment suggest the presence of small dormant populations of many species in complex microbiomes^8^. Even in nutrient-rich conditions such as laboratory batch cultures, steady state growth is ephemeral and unsustainable, because nutrients are rapidly depleted by growing heterotrophic bacteria^9^.

When colonising a finite niche, the depletion of essential nutrients is a common reason for bacteria to exit steady state growth. While the initial exit from exponential growth into stationary phase has been well-characterised in laboratory cultures, at least in *E. coli*^10–12^, ongoing biosynthesis and its regulation during protracted starvation is less explored. In part, this is historically due to technical challenges. Although imposing starvation for an essential nutrient in the laboratory can be straightforward, growth arrested cells severely reduce rates of key activities. Important regulatory changes to new protein synthesis can therefore have small and potentially undetectable impacts on measured total protein levels. Furthermore, diverse studies have suggested that population heterogeneity may increase in growth arrest^13,14^, such that the low average protein synthesis seen in these populations may result from a subset of the population that remains more active while other cells enter dormancy. Studying the low-level and heterogeneous activity maintained by growth arrested bacteria therefore requires highly sensitive methods to measure per-cell activity.

Fortunately, the development of methods for fluorescently labelling proteins, RNAs, and metabolites in single cells offers insight into the distribution of per-cell activity across non-growing populations. BONCAT (bio-orthogonal non-canonical amino acid tagging), is a powerful method to mark proteins newly synthesised during a specific period of labelling via incorporation of a click chemistry-compatible non-canonical amino acid (azidohomoalanine, AHA) by the native translational machinery^15,16^. BONCAT permits the estimation of total new protein synthesis per cell if the AHA in fixed cells is conjugated to a fluorescent dye^17^ and can also allow for enrichment and identification by LC-MS/MS of the nascent proteins made by the whole population of cells if lysate is conjugated with functionalised agarose beads^18^. Together, these methods can reveal how low levels of activity are distributed across growth-arrested populations and identify molecular players that comprise that activity.

A second obstacle to the study of growth arrest physiology and regulation is theoretical: in the absence of growth, the functions that should be prioritized for maximizing fitness are not immediately obvious. Previous work has suggested that even among Gammaproteobacteria, starvation survival strategies may be distinct, with some organisms retaining higher rates of new protein synthesis than others during protracted carbon starvation^17^. In many bacteria, starvation survival strategies include stockpiling non-limiting nutrients within high density stores, such as carbon-rich polyhydroxyalkanoate (PHA) granules^19^. Methods to evaluate mutant fitness, such as Tn-Seq/TRADIS, have permitted the identification of functionally important genes in diverse growth arrest conditions, allowing new insights into regulatory strategies that contribute to starvation survival^20,21^. By combining all these approaches, we hope to gain insight into the key functions that are preferentially maintained under severe resource limitation.

In this work, we have focused on the starvation responses of *Pseudomonas aeruginosa,* an opportunistic pathogen that has been shown to enter starved states in biofilms and other infection contexts^22–24^. We completely deprived planktonic cells of either a nitrogen source or a carbon source in defined minimal media for 24-96 hours as model growth-arrested conditions. Carbon starvation is expected to impose limitation for both energy and biosynthetic substrates (nucleotides and amino acids), while nitrogen starvation limits the availability of biosynthetic substrates but does not directly limit energy. In the absence of growth, there are no obvious constraints relating biosynthetic activity to cell size or ribosome abundance, so we have measured population size, culture optical density, cell size, ribosome abundance, and new protein biosynthesis to gain insight into global physiological parameters in these two different starvation conditions. We have also measured the total proteomes, newly synthesized (nascent) proteomes, and genes impacting fitness using standard label-free proteomics, BONCAT proteomics, and Tn-Seq. Finally, we investigated new protein synthesis and fitness determinants during transitions between starvations to explore conditions where remodelling the proteome might be beneficial despite ongoing growth inhibition.

We highlight three functional categories of genes that are relatively highly expressed during starvation and impact fitness: flagellar genes; proteases and chaperones; and the nitrogen-related PTS system (PTS^Ntr^). We show that mutations in representative genes from these categories cause diverse perturbations to total protein synthesis, ribosome abundance, morphology, and motility during growth arrest. When considering all our data, we propose that starved *P. aeruginosa* is able to store non-limiting nutrients and dynamically redistribute these resources to power key activities. Ultimately, we hope that these datasets can serve as a resource for further explorations of physiology in starvation-induced growth arrest.

## Results

### Defining the translational capacity of growth-arrested *P. aeruginosa*

We first broadly characterised the response of *P. aeruginosa* to complete starvation for nitrogen or carbon sources. Overnight *P. aeruginosa* LB cultures were pelleted and resuspended at low density (OD_500_ 0.2) in MOPS-buffered minimal media lacking either ammonium chloride (nitrogen source) or sodium succinate (carbon source) and were sampled across several days. As expected, the absorbance of starved cultures did not measurably increase within the timeframe of typical bacterial growth curves (Supplemental Figure 1A), suggesting that cells did not grow under either starvation condition. However, clear distinctions in culture absorbance, CFU counts, and cell area became apparent when starved cultures were tracked for prolonged periods. Carbon starved cultures maintained relatively stable absorbance and CFU counts with prolonged incubation, while average cell area as a measure of cell size steadily decreased (Figure 1A-C), suggesting a gradual loss of biomass. Nitrogen starved cultures showed more dynamic trends, increasing in CFU counts during the first 8 hours of starvation before stabilizing (Figure 1A). The absorbance of nitrogen starved cultures almost doubled over 100 hours of incubation (Figure 1B) and the average area of nitrogen starved cells quickly decreased following resuspension in starvation media before increasing with prolonged incubation (Figure 1C). These trends could be explained by initial reductive divisions upon resuspension in nitrogen starvation media, as has been observed in *E. coli*^25^, followed by subsequent changes in cell morphology and culture opacity that occur independently of cell divisions. Interestingly, motile cells were frequently observed in both nitrogen starved and carbon starved cultures (Figure 1D), suggesting that flagellar motility is maintained by a sub-population of non-growing *P. aeruginosa* during days of nutritional deprivation, an observation also recently reported^26^. In general, *P. aeruginosa* showed dynamic responses to the distinct starvations and, following an initial ∼10-hour adaptive phase, cells appeared to enter a prolonged growth-arrested state with stable population numbers in both starvation conditions.

We then probed the translational capacity of growth-arrested *P. aeruginosa* under both starvation conditions. BONCAT was used as a proxy to measure ongoing protein synthesis in starved cells. Starved cultures were amended with azidohomoalanine (AHA) for various labelling periods before cells were fixed and a fluorophore was covalently linked to incorporated AHA by strain-promoted azide-alkyne cycloaddition. Azidohomoalanine does not appear to be catabolized by *P. aeruginosa* and was unable to alleviate carbon or nitrogen limitation to permit growth (Supplemental Figure 1B, C). Although the average signal per starved cell was much lower than that observed for growing cells (Supplemental Figure 1E, G), it increased approximately linearly with labelling time in both starvation conditions (Figure 1E, F). Fluorescence distributions did not show evidence of distinct sub-populations. Although AHA incorporation is not an absolute measure of protein synthesis, nitrogen-starved cells consistently showed higher signal per labelling time than carbon-starved cells (Figure 1G, H), indicating relatively higher rates of new protein synthesis.

The ribosome content of growth arrested *P. aeruginosa* was then probed under both starvation conditions using fluorescence in-situ hybridisation (FISH)^6,27^. Starved cells showed lower signal than growing cells (Supplemental Figure 1D, F), and nitrogen starved cells had lower signal on average than carbon starved cells despite the BONCAT indications of higher protein synthesis rates (Figure 1I). Furthermore, while nitrogen starved cells maintained a low and stable average ribosome count across days of starvation, the higher average signal of carbon starved cells waned with time (Figure 1I). These data indicated that distinct and dynamic translational strategies accompany the responses to the two starvation conditions, and that ribosome content and protein synthetic rate are not directly correlated in growth-arrested *P. aeruginosa*.

**Figure 1.**
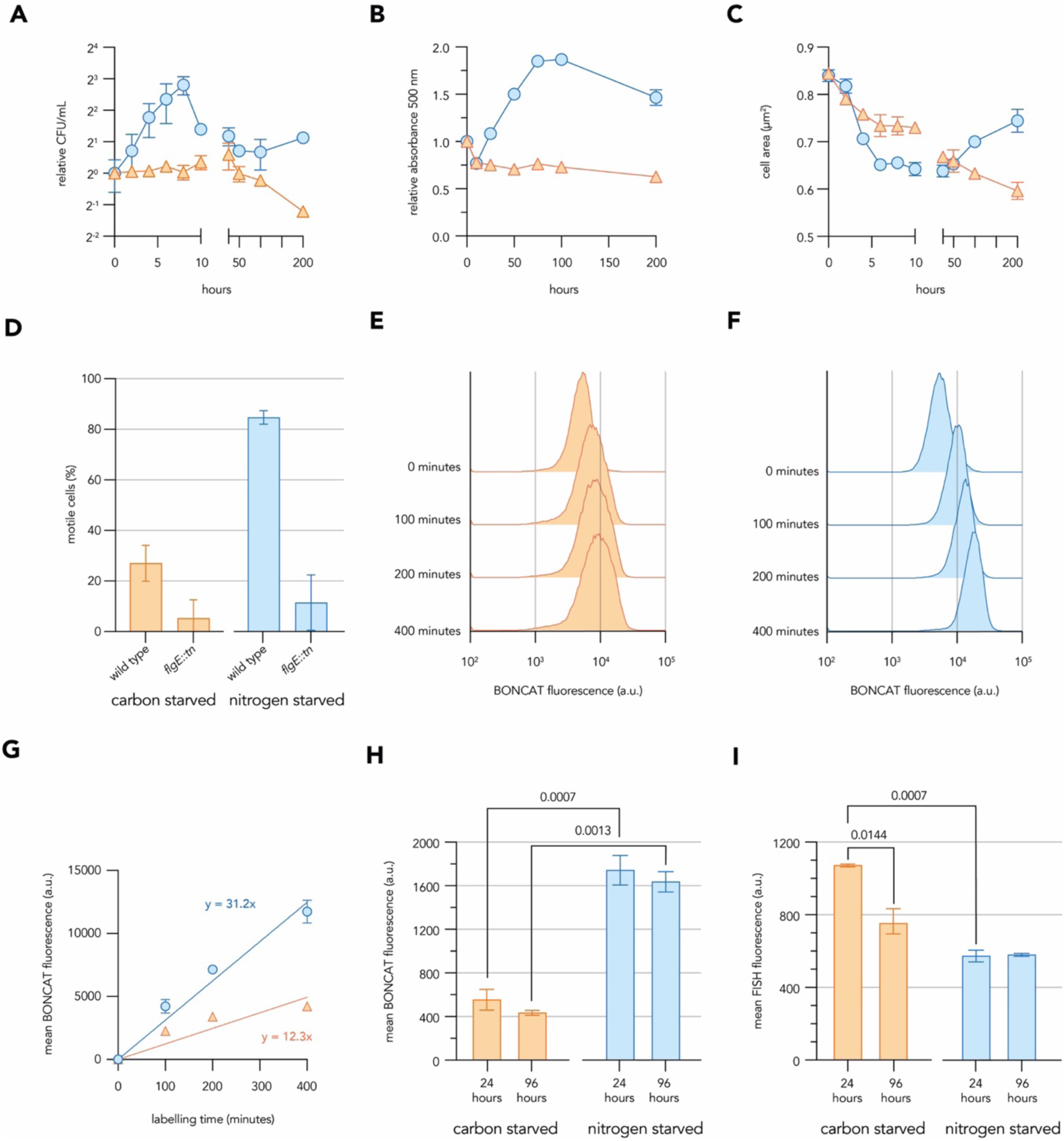
Defining the protein synthetic capacity of growth-arrested *P. aeruginosa*. Longitudinal trends in colony-forming units (**A**); absorbance 500nm (**B**); and average cell area (**C**) of *P. aeruginosa* cultures following resuspension in nitrogen starvation minimal media (blue circles) or carbon starvation minimal media (orange triangles). Cell area measurements were derived from the microscopy of samples fixed at indicated timepoints. Bar graphs (**D**) show the percentage of motile cells quantified from videos of *P. aeruginosa* cultures sampled following 48 hours of starvation for carbon or nitrogen in minimal media. Histograms show the distributions of new protein synthesis levels as per-cell fluorescence values in *P. aeruginosa* cultures starved of carbon (**E**) or nitrogen (**F**) in minimal media for 48 hours and BONCAT labelled with 500 µM AHA for time periods indicated. Line graph (**G**) show the relationship between BONCAT labelling time and average fluorescence of *P. aeruginosa* cells starved of carbon (orange triangles) or nitrogen (blue circles) in minimal media for 48 hours, with the gradient of each line shown. Mean fluorescence values of *P. aeruginosa* cells starved of carbon or nitrogen in minimal media for 24 or 96 hours and BONCAT labelled for 200 minutes with 500 µM AHA (**H**) or FISH stained with ribosome-directed probes (**I**). All data represent the mean value calculated from at least three biological replicates with standard deviation displayed as error bars. Histograms are representative of at least three biological replicates. T-tests were used to calculate the p-values denoted above select comparisons, with only significant values (p < 0.05) being shown. a.u. = arbitrary units.

### Starved *P. aeruginosa* undergo bursts of translation during a starvation transition

Having defined two growth-arrested states with distinct translational activities, we next investigated how starved *P. aeruginosa* reacted to rapid transitions between the two. We collected nitrogen-starved cells by centrifugation and resuspended them in carbon starvation medium, hypothesising that quickly shifting between starvation conditions would force cells to adapt their activities despite ongoing growth-arresting nutrient limitation. No substantial increase in CFU counts or culture absorbance followed the transition from nitrogen starvation to carbon starvation media, indicating that the media switch did not leave sufficient residual nutrients to support growth (Supplemental Figure 2A, B).

Two-hour BONCAT labelling pulses were used to compare new protein synthesis before, during and after the transition from nitrogen starvation to carbon starvation media (Figure 2A) and a control transition into fresh nitrogen starvation conditions (Figure 2D). A large burst of translation accompanied the transition of cells from nitrogen starvation to carbon starvation conditions, with the average BONCAT signal increasing 3-fold immediately following the transition and subsequently decreasing (Figure 2B, C). Within a day of the transition the average BONCAT signal decreased to the low levels previously observed during carbon starvation (Figure 1G. H). These translational bursts were also observed in transitions from carbon to nitrogen starvation medium (Supplemental Figure 2F, G, H), although in this case an upshift in protein biosynthesis could be predicted by the higher average activity maintained during nitrogen starvation. Transitions from nitrogen starvation medium into fresh nitrogen starvation media elicited only a slight increase in average BONCAT signal (Figure 2E, F), possibly due to replenishment of the carbon source, which should be consumed during ongoing nitrogen starvation. These data highlight that growth-arrested *P. aeruginosa* can quickly upregulate biosynthetic activity in response to environmental changes even during prolonged starvation, indicating that cells may maintain latent biosynthetic capacity during nutritional deprivation.

**Figure 2.**
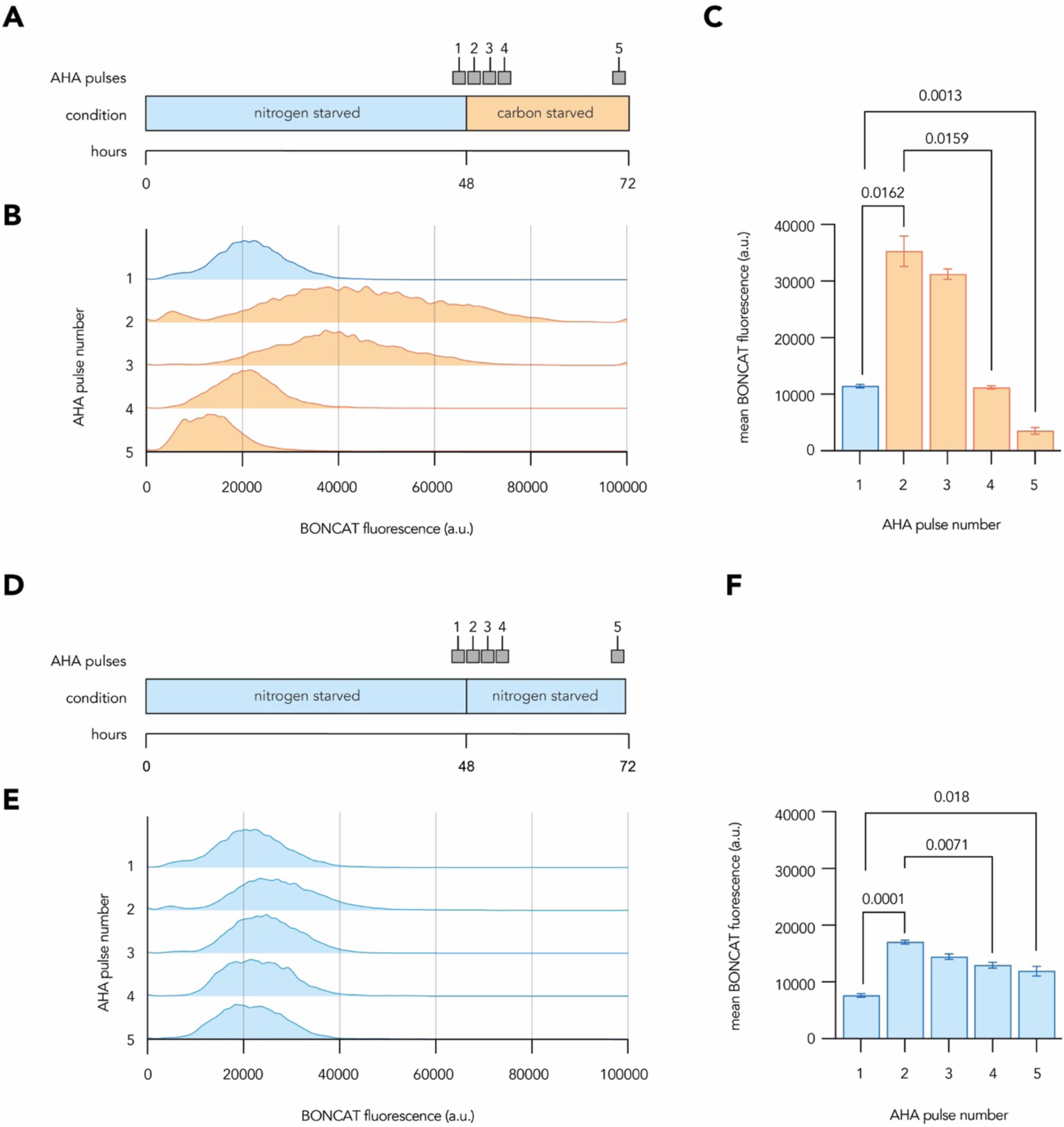
Starved *P. aeruginosa* undergo bursts of translation during a nutritional transition. Schematics show the time-course of two-hour 500 µM AHA pulses applied to *P. aeruginosa* cultures for BONCAT labelling during a transition from nitrogen starvation to carbon starvation (**A**) and the transition from nitrogen starvation to new nitrogen starvation minimal media (**D**). Histograms (**B** and **E**) show the distributions of per-cell fluorescence values from fixed BONCAT samples labelled across a transition between nitrogen and carbon starvation. Bar graphs (**C** and **F**) show the mean BONCAT fluorescence values retrieved for each AHA labelling pulse. All data represent the mean value calculated from at least three biological replicates with standard deviation displayed as error bars. Histograms are representative of at least three biological replicates. T-tests were used to calculate the p-values denoted above select comparisons, with only significant values (p < 0.05) being shown. a.u. = arbitrary units.

### The total and nascent proteomes of growth-arrested *P. aeruginosa*

We next investigated the portfolio of proteins present in and being produced by growth-arrested *P. aeruginosa* during nitrogen starvation, carbon starvation and when transitioning from nitrogen to carbon starvation. We first applied label-free quantification to the total proteome of *P. aeruginosa* by LC-MS/MS at several key points during a starvation time course including 48 hours in nitrogen starvation, 24 hours in carbon starvation, and a final outgrowth in nutrient-replete minimal medium allowing sampling during exponential phase (Figure 3A). We normalized quantities of each protein to the total summed abundances of all proteins in each sample and identified more than 3900 proteins with an average abundance of more than 1 part per million (ppm) in each condition. Principle component analysis demonstrated a tight clustering of replicate proteomes and highlighted that the proteomes of exponential phase *P. aeruginosa* cultures were distant from those of starved or transitioning cultures along principal component 1, which accounted for 61.1% of the variation (Figure 3B). When the hundred most abundant proteins from total proteomes were compared across conditions we found substantial overlap, particularly among the most abundant proteins identified from starved and transitioning cultures, with 46 abundant proteins being shared among all nutrient-limited conditions (Figure 3C, Supplemental File 1). These abundant proteins were then functionally classified using a modified PseudoCAP^28^ scheme (Supplemental File 1) to allow for comparison of major cellular priorities under each condition. The abundant proteins of exponential phase proteomes were particularly enriched in translation-associated functions, while proteins relating to core cellular functions such as carbon compound anabolism and energy metabolism were abundant in all proteomes. (Supplemental Figure 3A).

**Figure 3.**
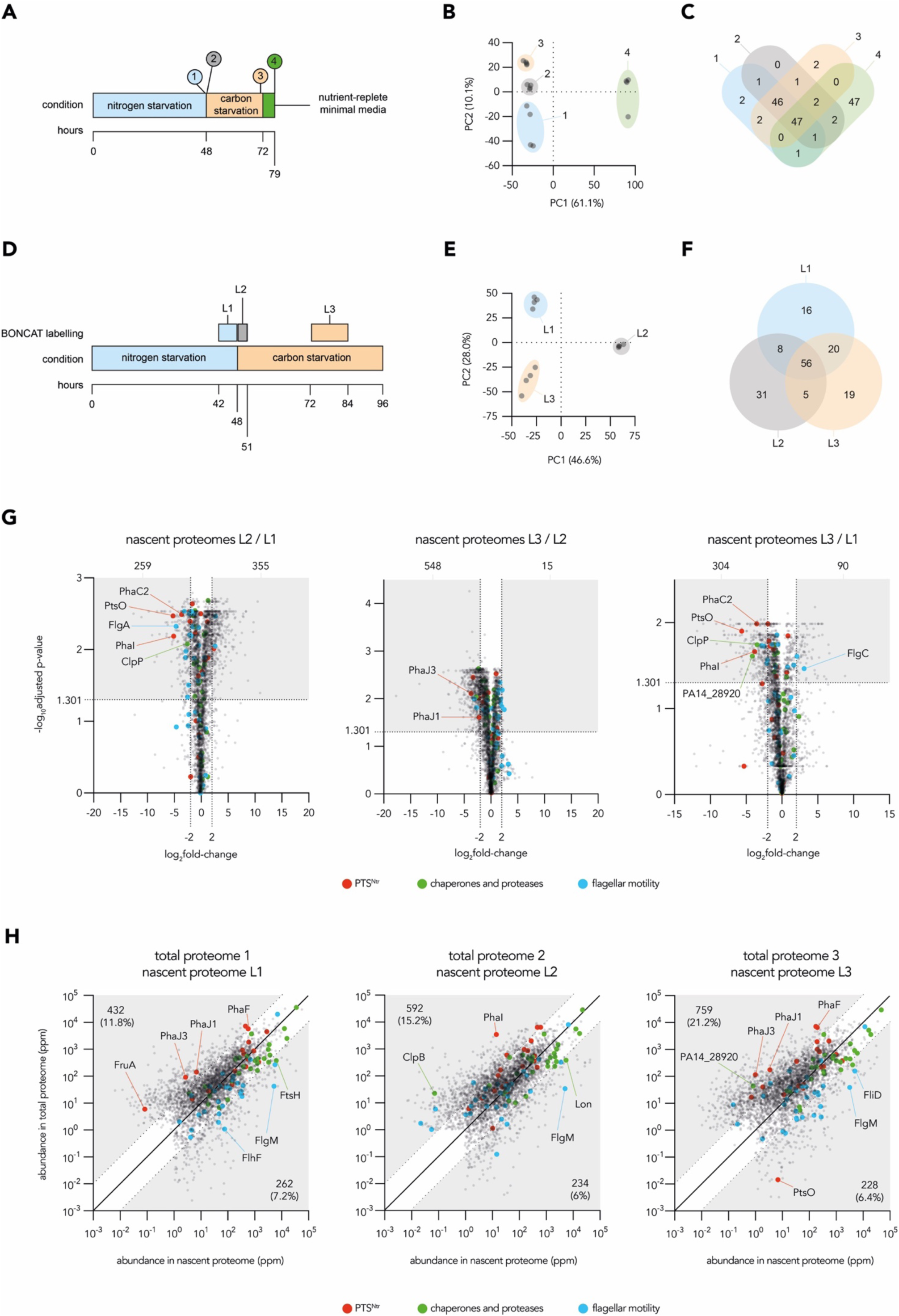
The total and nascent proteomes of growth-arrested *P. aeruginosa*: Schematics show the time-courses employed in total proteomic (**A)** and nascent (BONCAT) proteomic (**D**) experiments. Scatter plots show principal component analyses of returned total (**B**) or BONCAT (**E**) proteomes. Venn diagrams show the overlap of the top 100 most abundant proteins from each timepoint of total proteomic (**C**) and BONCAT proteomic (**F**) experiments. Volcano plots (**G**) show the fold-change in abundance of nascent proteins when comparing different time points of the BONCAT proteomic experiment. In each volcano plot, shaded boxes highlight the nascent proteins which passed fold-change (log_2_foldchange > 2 or < -2) and adjusted p-value (log_10_adjusted-p-value > 1.301) cut-offs to be considered as robust changes in abundance between compared proteomes, with the number of such proteins denoted above each shaded box. Scatter plots (**H**) combine total and nascent proteomic data and show the normalised average abundances of individual proteins detected in both datasets. Shaded regions indicate proteins that had greater than 10-fold difference in relative abundance between the two types of proteomic experiments, and the fraction of all detected proteins that fall within each shaded region is indicated. The larger coloured dots highlight individual genes belonging to key physiological functions. All analyses used mean abundances calculated from four biological replicates.

Relatively few distinctions were noted when comparing the abundance of individual proteins between the proteomes of starved and transitioning cultures, likely due to low amounts of new protein synthesis being overshadowed by the abundant proteome established at the onset of growth arrest^29^ (Supplemental Figure 3B). Therefore, we employed BONCAT-linked proteomics to enrich and quantify the newly synthesized proteomes of *P. aeruginosa* under our conditions of interest. Cultures were starved of nitrogen for 48 hours, the last six of which were labelled with AHA; then transitioned to carbon starvation, with a three-hour labelling period commencing immediately after the media switch, and a second labelling period of 12 hours started after 24 hours of carbon starvation (Figure 3D). The labelling times were chosen to compensate for different rates of per-cell new protein synthesis observed in our previous fluorescence experiments (Figure 2A, B, C). AHA-labelled proteins were enriched and analyzed by LC-MS/MS, as for the total proteomes. We identified more than 3200 proteins with an average abundance of more than 1 ppm in each nascent proteome. Principle component analysis of the nascent proteomes revealed a tight clustering of replicates but suggested that the portfolios of nascent proteins made during different labelling periods were distinct (Figure 3E). Indeed, the hundred most abundant proteins from each nascent proteome overlapped considerably less than the most abundant proteins in the total proteomes (Figure 3F, 3C). Interestingly, we found similar patterns of functional enrichment among the nascent proteins from each condition, suggesting that although *P. aeruginosa* may express distinct proteins during each labelling period, similar general functions are maintained throughout the three labelling periods investigated (Supplemental Figure 3C).

When directly comparing nascent proteomes, we found that the abundance of hundreds of proteins differed substantially and significantly between the three labelling periods (Figure 3G). Proteins from three functional categories of particular interest are highlighted: proteases and chaperones; flagellar motility; and the PTS^Ntr^ system and its regulon (Figure 3G, Supplemental Figure 3B; groups defined in Supplemental File 1). These categories show relatively high expression, interesting dynamics, and/or functional importance in Tn-Seq datasets (as described below). As similar time-courses were used to generate the total and nascent proteomic datasets, we compared the relative abundances of individual proteins detected at analogous time points using scatter plots (Figure 3H, Supplemental File 1). In general, new protein synthesis reflects the composition of the total proteome, consistent with ongoing maintenance to broadly preserve cellular composition. However, points deviating from the diagonal reflect active proteome remodelling or proteins with unusually high or low turnover rates. Interestingly, many components of the PTS^Ntr^ network, such as proteins involved in the synthesis of polyhydroxyalkanoate (PHA) granules^30^, have higher relative abundance in the total proteomic data, perhaps consistent with expression during the entry to growth arrest. Many flagellar proteins, proteases, and chaperones are represented more strongly in the nascent proteomes, indicating ongoing enhancement of their levels or high turnover. Overall, these data provide a detailed and comprehensive catalog of the total and nascent proteins in starved *P. aeruginosa*.

### The fitness determinants of growth-arrested *P. aeruginosa*

We next asked which genetic factors contributed to survival during our starvation conditions and across transitions between them. We utilized a high-density transposon mutant library to perform transposon-insertion sequencing and identify mutants that significantly changed in abundance after exposure to nitrogen or carbon starvation conditions. We also investigated how abundances changed following a transition to a second starvation condition (Figure 4A). For each starvation sample collected, a short (approx. 2 hrs, 3 doublings) outgrowth in LB medium was performed to allow starvation-imposed fitness defects to be amplified.

While many genes affected fitness during the first starvation exposure, only a small number of genes specifically impacted fitness across a starvation transition (Figure 4B, Supplemental file 1). Interestingly, many genes impacted fitness similarly in both starvation conditions (Figure 4C/D), hinting that conserved processes and pathways generally support viability during growth-arrest. Many genes highlighted in the nascent proteomic data robustly decreased in transposon read counts during carbon and/or nitrogen starvation, suggesting that expressing these proteins was beneficial for *P. aeruginosa* survival during prolonged growth arrest (Figure 4E, Supplemental file 1). All genes whose transposon read counts significantly and substantially changed during either starvation condition were then functionally categorized (Figure 4F). We found that survival-enhancing and survival-diminishing genes identified from analyses of either starvation condition were similarly distributed across functional categories. Interestingly, many genes encoding flagellar components increased in transposon read counts specifically following the transition of nitrogen starved cultures to carbon starvation conditions (Figure 4E). These data suggest that flagellar motility is costly, and while investing in exploration may have benefits in natural environments, loss of motility in the laboratory can improve fitness of growth-arrested *P. aeruginosa* navigating this transition. Overall, transposon insertion sequencing provided a holistic view of the fitness determinants impacting *P. aeruginosa* survival during starvation. In combination with corresponding proteomics datasets, these fitness data permit the identification of pathways and processes which are that prioritised and impactful during dynamic growth arrest.

**Figure 4.**
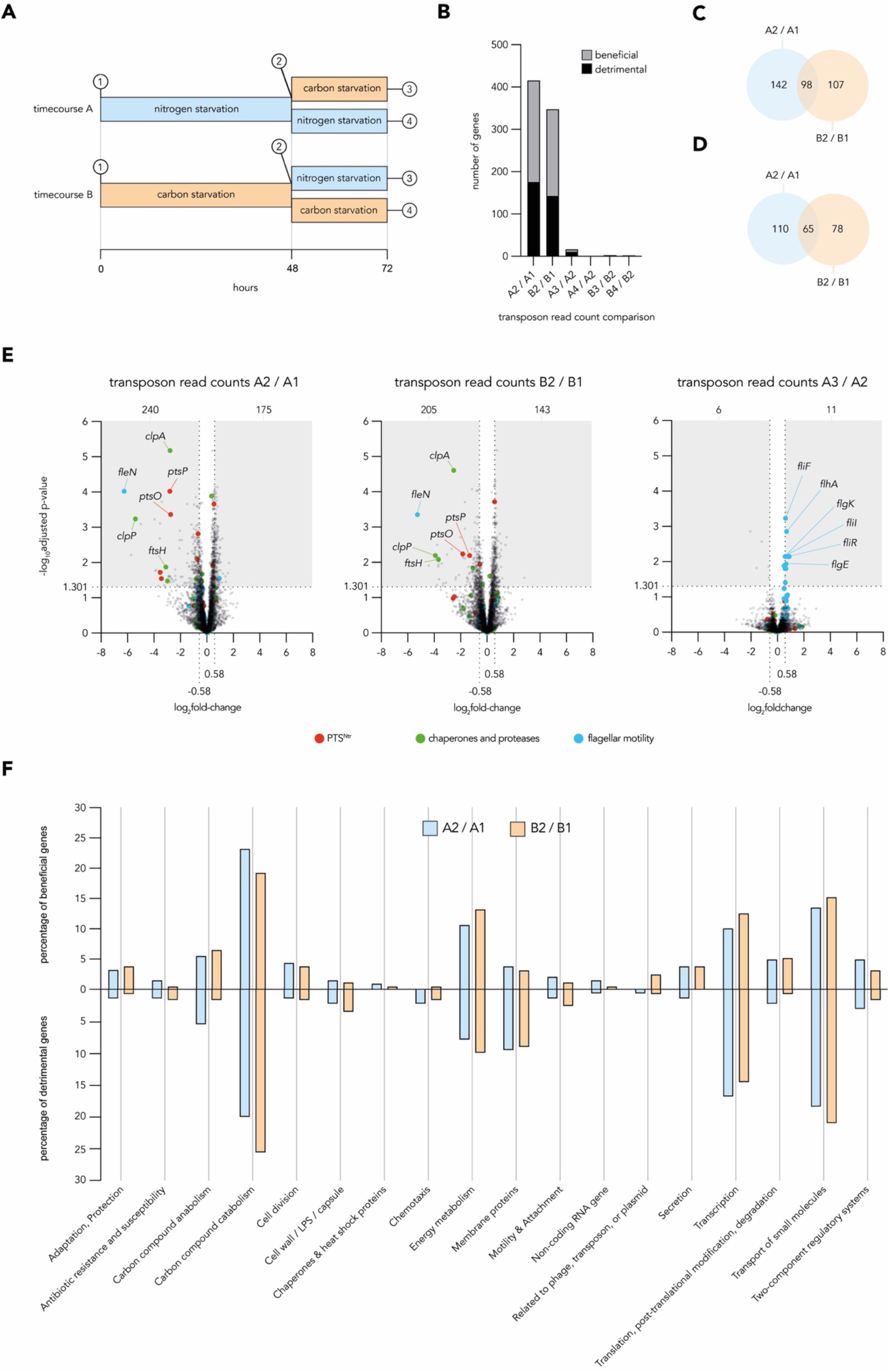
The fitness determinants of growth-arrested *P. aeruginosa*: Schematic (**A**) shows TnSeq experimental time courses, highlighting the timepoints sampled and used in subsequent analyses. Bar graph (**B**) shows the numbers of beneficial genes whose transposon read count robustly decreased and detrimental genes whose transposon read count robustly increased in comparisons of library read counts between sampled timepoints. Fitness changes were considered robust if read counts crossed fold-change (log2fold-change > 0.58 or < -0.58) and adjusted p-value (log_10_adjusted-p-value > 1.301) cut-offs when comparing samples. Venn diagrams (**C**) show the overlap of robust beneficial genes and robust detrimental genes (**D**) in comparisons relevant to survival during nitrogen starvation (top) and carbon starvation (bottom). Volcano plots (**E**) show the fold-change in transposon read counts and adjusted p-values for genes when comparing different sampling periods. In each plot, shaded boxes highlight the genes whose read count passed fold-change and adjusted p-value cut-offs to be considered robust, with the number of these robust hits denoted above each shaded box. The larger coloured dots highlight individual genes belonging to key physiological functions. Bar graphs (**F**) show the percentage occupancy of modified PseudoCAP groups by robust beneficial genes (top) and robust detrimental genes (bottom) retrieved from the comparisons of library read counts for nitrogen starved or carbon starved samples. All analyses used mean read counts calculated from at least five biological replicates.

### Patterns of expression and fitness impacts for key physiological functions

Having generated proteomic and TnSeq datasets from growth-arrested *P. aeruginosa*, we next concentrated on three functional categories that were generally highly expressed and important for fitness: flagellar motility; proteases and chaperones; and the PTS^Ntr^ phosphotransferase system (Supplemental Figure 4A). We inspected the nascent proteome abundances and Tn-Seq read counts of individual components to analyze functional relationships within these categories (Figure 5).

**Figure 5.**
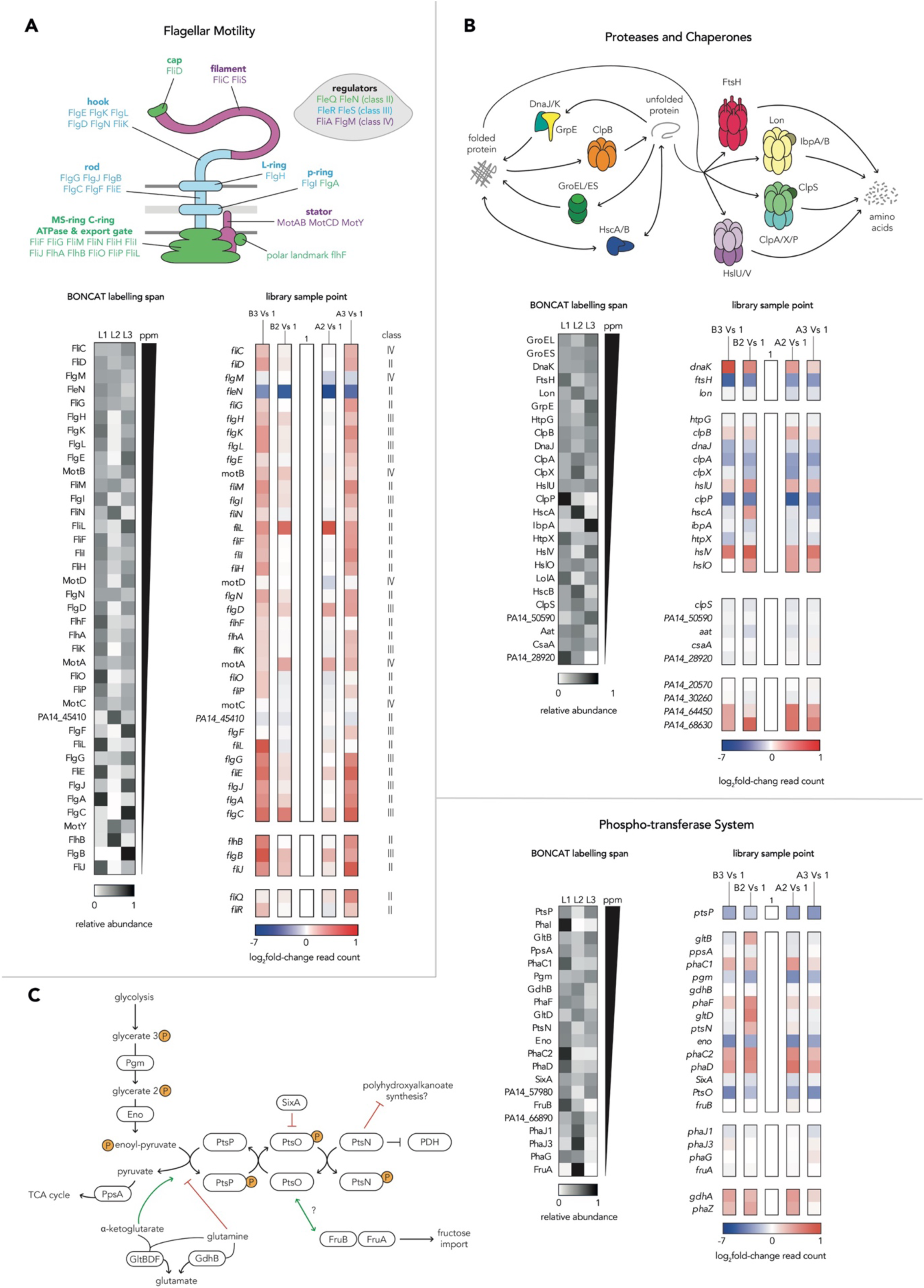
Patterns of expression and read counts for key physiological functions: Schematics portray the structural or metabolic interplay of components belonging to each of the three key physiological functions highlighted by proteomic and TnSeq data. Genes and proteins of interest are grouped into three key pathways: flagellar motility (**A**), proteases and chaperones (**B**) and the phospho-transferase system (**C**). Black and white heat maps portray the fraction of total abundance, summed across the three labelling periods of the BONCAT proteomics experiment, associated with each time point. In each map, nascent proteins are ordered top to bottom by the total abundance summed across all three labelling periods. Coloured heat maps portray the log2 fold-change in transposon read counts of genes of interest when comparing sampling timepoints of the transposon-insertion sequencing experiment. Each comparison of starved (A2 and B2) or transitioned (A3 and B3) read counts are made relative to stationary phase read counts (A1 and B1). Gaps in heat maps indicate missing data where nascent protein was not detected or a gene was absent from the transposon-mutant library. All analyses of BONCAT proteomic abundances used the mean abundances calculated from four biological replicates and analyses of TnSeq read counts used the mean read counts calculated from at least five biological replicates.

Collectively, flagellar proteins constitute roughly 2% of the nascent protein retrieved from all three BONCAT labelling timepoints, although the flagellin protein FliC always ranked amongst the top 25 most abundant individual nascent proteins (Supplemental Figure 4B). Nascent flagellar proteins are most abundant during nitrogen starvation conditions and drop in relative abundance in the nascent proteomes of transitioning and carbon starved cells (Figure 5A). Some highly abundant flagellar proteins and regulators (FliC, FliD, FlgM, FleN and FlgG) were expressed during all three labelling timepoints, but expression of many other flagellar components was dynamic. In general, proteins expressed earlier in flagellar biosynthesis, as part of the “class II” set transcriptionally regulated by FleQ, were more likely to have their highest relative expression during nitrogen starvation, while some members of the later classes “III/IV” regulated by FleR and FliA had notable bursts of new synthesis after the switch to carbon starvation^31^. During individual starvations, read counts for most flagellar structural genes remain stable, implying flagellar defects had minimal impact to fitness during stable growth-arrested conditions. In contrast, read counts from genes encoding negative transcriptional regulators of flagellar biosynthesis, *fleN* and *flgM,* sharply decreased during both starvations, suggesting a high fitness cost of dysregulated overexpression of flagellar components. Together with the increased read counts for flagellar structural genes observed during starvation transitions, these data suggest that growth arrested *P. aeruginosa* carefully regulate ongoing flagellar activity. Nevertheless, proteomics data indicate that new synthesis of flagellar components remains a priority throughout prolonged starvation for *P. aeruginosa*.

Proteases and chaperones were extremely abundant in the nascent proteomes of growth-arrested *P. aeruginosa*, with this group constituting roughly 10% of nascent proteins at each labelled timepoint (Supplemental Figure 4B). Several individual proteases and chaperones, including GroEL, GroES, DnaK, FtsH, and Lon, were constitutively expressed and among the most abundant nascent proteins retrieved from all three labelling periods (Figure 5B, Figure 3). Many non-essential protease and chaperone genes, such as *clpA*, *clpX, clpP*, and *ftsH*, exhibited strong and significant decreases in their read counts during starvation, suggesting components of this group are crucial for survival of growth arrest. Although read count increases were also observed in some proteases (for example, *hslUV, dnaK*), these were small in magnitude and failed to pass thresholds of significance (Supplemental file 1). For most proteases and chaperones, similar fitness impacts were observed regardless of the starvation condition, suggesting that they globally and generally contribute to starvation survival.

Transposon mutants of *ptsP* and *ptsO*, the first two components of the PTS^Ntr^ phosphorylation cascade, had some of the strongest and most significant survival defects of any genes during both carbon and nitrogen starvations (Figure 4E, Figure 5C, Supplemental File 1). This led us to interrogate the complex and enigmatic regulatory network surrounding the PTS^Ntr^ in *P. aeruginosa*. PTS^Ntr^ is highly conserved and modulates fluxes of carbon and nitrogen through bacterial metabolism via phosphorylation of PtsN, which has been proposed to regulate activity of diverse interaction partners in its unphosphorylated state^32–34^. The phosphate derives from phosphoenolpyruvate, a central carbon metabolite, and PtsP activity is affected by 2-ketoglutarate and glutamine, providing a connection to nitrogen metabolism^35^ (Figure 5C). We interrogated our expression and fitness data for components of the PTS^Ntr^ and fructose-related PTS systems; for metabolic enzymes impacting phosphoenolpyruvate, 2-ketoglutarate, and glutamine; and for known regulators of PHA metabolism, which is reported to be impacted by PTS^Ntr^ in Pseudomonads^36^. Transposon sequencing data suggested that in addition to *ptsP* and *ptsO,* genes required for phosphoenolpyruvate synthesis (*eno* and *pgm*) were important for survival during both starvation conditions (Figure 5C). PTS^Ntr^-related proteins constitute less than 1% of the nascent protein recovered from BONCAT labelling at each timepoint (Supplemental Figure 4B). However, some of the most highly produced proteins in this group (PtsP, PhaI, PhaC1) showed dynamic changes to expression, with strong decreases in abundance during the transition from nitrogen starvation to carbon starvation (Figure 3G, 5C). In contrast, proteins with putative roles in breakdown of PHA (PhaJ1, PhaJ3) increased during the transition.

### Validating Impacts of Three Key Functions in Growth-Arrested *P. aeruginosa*

Finally, we returned to our measures of per-cell size, ribosome abundance, and protein synthesis to investigate whether mutations in key representative genes identified from global proteomic and TnSeq datasets impacted these measures of growth arrested physiology. Based on observations from our global datasets that PTS-regulated processes were important, we also investigated PHA production using BODIPY^493/503^ staining^37^.

To inspect the role of proteases and chaperones during growth arrest we focused on *ftsH* as a representative protease^20^. We found that *ΔftsH* cells had severely dysregulated new protein synthesis, producing on average substantially more protein during nitrogen starvation and substantially less during carbon starvation relative to wild type cells (Figure 6A). Concurrently, these Δ*ftsH* cells on average possessed more ribosomes in nitrogen starvation and fewer ribosomes in carbon starvation relative to wild type cells (Figure 6B) and failed to produce a burst of protein synthesis when transitioned between starvation media (Figure 6C). These data complement previous findings that *ftsH* contributes to fitness during growth arrest in *P. aeruginosa* by degrading damaged proteins, thus limiting the accumulation of toxic aggregates^38^. Indeed, Δ*ftsH* exhibited a slight fitness defect when directly competed against wild-type cells during numerous starvation transitions (Figure 6D). These data suggest that individual proteases can dramatically, differentially and consequentially sculpt the biosynthetic capacity of growth-arrested *P. aeruginosa* during nitrogen and carbon starvation.

**Figure 6.**
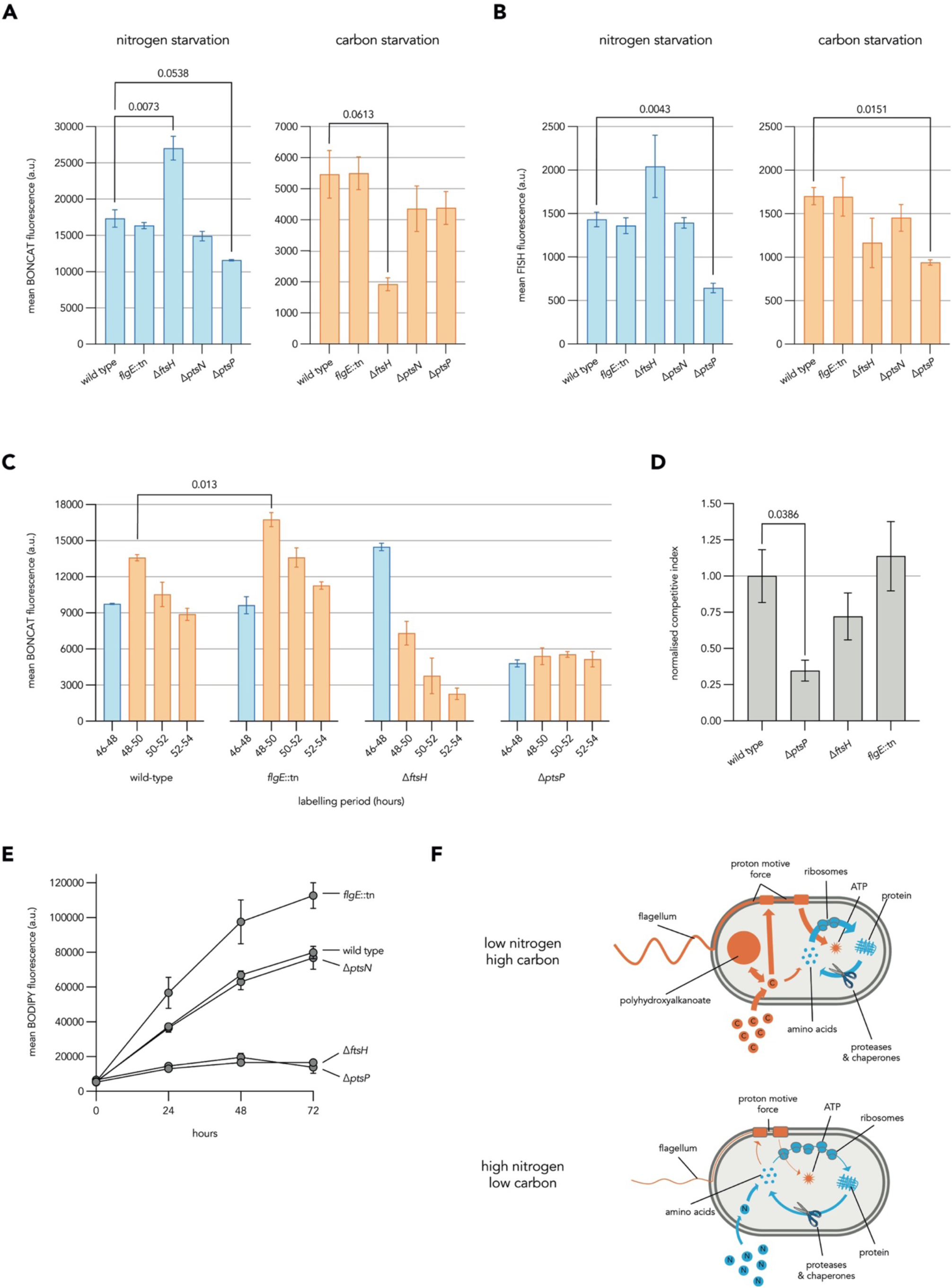
Validating Impact of Three Key Features in Growth-Arrested *P. aeruginosa* Bar graphs. (**A**) show the average cellular BONCAT fluorescence from the labelling of *P. aeruginosa* cultures for 240 minutes with 200 µM AHA following 48 hours of incubation in nitrogen starvation (left) or carbon starvation (right) minimal media. Bar graphs (**B**) show the average cellular FISH fluorescence from the staining of *P. aeruginosa* cultures following 48 hours of incubation in nitrogen starvation (left) or carbon starvation (right) minimal media. Bar graphs (**C**) show the average cellular fluorescence from pulsed BONCAT labelling of starved *P. aeruginosa* cultures during a transition between nitrogen starvation (blue) and carbon starvation (orange) minimal media. Cultures were incubated in nitrogen starvation minimal media for 48 hours prior to the transition, with two-hour 1000 µM AHA labelling pulses applied prior to (46-48) or following (48-50, 50-52, 52-54) the transition. Bar graph (**D**) shows the normalised competitive index of strains competed against wild-type *P. aeruginosa* during three transitions between nitrogen starvation and carbon starvation minimal media (**D**). Line graphs (**E**) show PHA content as average per-cell BODIPY fluorescence of *P. aeruginosa* populations fixed at indicated timepoints during nitrogen starvation. Schematics (**F**) highlight the interplay of prioritised cellular processes and relevant resource stores in *P. aeruginosa* during growth arrest under different starvation conditions. Arrow and line thickness represent the contribution of processes to the use of carbon or nitrogen sources in either starvation situation, as inferred from experimental data. All data represent the mean value calculated from at least three biological replicates with standard deviation displayed as error bars. T-tests were used to calculate the p-values denoted above select comparisons, with only values close to or passing significance thresholds (p < 0.05) being shown. a.u. = arbitrary units.

We used non-motile transposon insertion mutant *flgE::MAR2xT7*^39^ to probe the influence of flagellar motility during growth arrest (Figure 1D). Although no substantial change to protein synthesis and ribosome counts were found during carbon or nitrogen starvation, *flgE::tn* cells had a significantly larger burst of protein synthesis when transitioned between starvation conditions (Figure 6A, B, C). Interestingly, these cells also stored more PHA than wild type cells during nitrogen starvation (Figure 6E). Together, these observations suggest that mutational loss of the capacity for flagellar motility frees resources for other biosynthetic activities during starvation.

Deletion mutants Δ*ptsP*^20^ and Δ*ptsN* were used to probe impacts of the PTS during growth arrest. While the growth of Δ*ptsP* in nutrient-rich conditions was similar to wild type, Δ*ptsN* exhibited a growth defect, suggesting that unphosphorylated PtsN may play important roles during growth, while its phosphorylation is required for fitness during growth arrest (Supplemental Figure 5A). Both mutants had slightly reduced protein synthesis relative to wild type during nitrogen and carbon starvation (Figure 6A), with Δ*ptsP* cells also possessing significantly fewer ribosomes during both starvation conditions (Figure 6B). This mutant also failed to produce a translational burst during transitions between starvation conditions and exhibited a strong fitness defect when competed against wild type cells during transitions (Figure 6C, D). Furthermore, Δ*ptsP* cells appeared to elongate and failed to accumulate PHA during nitrogen starvation, as reported previously for other organisms^36^ (Figure 6E, Supplemental Figure 5C). These data show that the PTS^Ntr^ impacts morphology and biosynthetic capacity in starved *P. aeruginosa* and validate *ptsP* as a crucial regulator of physiology in these growth-arrested contexts.

Overall, we attributed significant defects to the disruption of individual components of each of the three key functional categories identified by proteomic and TnSeq data. Proteases and chaperones and the PTS^Ntr^ appear to severely and pleiotropically impact the physiology of starved cells, while the disruption of flagellar motility was specifically consequential during transitions, as suggested by the TnSeq data (Figure 5). Interestingly, defects in PHA accumulation and ribosome abundance were observed for mutants with strong fitness defects in both nitrogen and carbon starvation conditions.

## Discussion

By combining quantitative single-cell techniques, BONCAT proteomics, and transposon-insertion sequencing we probed the physiology of starved *P. aeruginosa* cells. Taken together, our results show that starvation-induced growth arrest is dynamic and actively regulated, with diverse new proteins continuously being synthesized for days by most cells in the population, albeit at much lower rates than during exponential growth. Our data offer insights into strategies that might support these dynamics. Growth arrest due to absence of an essential macronutrient provides opportunities for storing other resources available in excess. Under fluctuating growth-arrested conditions, such internal stores can contribute to ongoing biosynthesis. Regulatory strategies must then determine when and how stored resources should be expended to meet challenges imposed by the environment. We propose three example intracellular resource storage mechanisms, relevant under different conditions of limiting and excess nutrients: ribosomes and other cellular proteins as storage for excess nitrogen, PHA granules as storage for excess carbon, and the proton motive force (PMF) as a store of energy (Figure 6F). Resources can be shifted between these stores during starvation-induced growth arrest, and the three functional pathways we have highlighted (proteases and chaperones, flagella, and the PTS^Ntr^ system) are involved in driving and regulating these shifts. Ribosomes and proteins represent a major internal store of amino acids that could be liberated in times of need^40^. During nitrogen starvation, where energy is available but amino acids are limiting for protein biosynthesis, it may be advantageous to more quickly recycle ribosomes and proteins to salvage nitrogen^40,41^ than during carbon starvation. Many bacterial species store ribosomes in inactive states during starvation by binding ribosome hibernation factors^42–44^ and translation elongation can be slowed to different degrees in response to distinct nutrient limitations^45,46^. We observed evidence of slowed or hibernating ribosomes during carbon starvation, where cells showed relatively low protein synthesis, but high ribosome abundance compared to nitrogen starvation (Figure 1H, I). Interestingly, the deletion of the protease *ftsH* leads to major dysregulation of ribosome abundance, new protein synthesis and cell morphology during starvation (Figure 6A-C, Supplemental Figure 5B). The increase in ribosome abundance during nitrogen starvation in Δ*ftsH* cells relative to the wild type is potentially consistent with a decrease in ribosome turnover. However, this mutant was also defective in maintaining ribosomes during carbon starvation, suggesting a more complex role of the protease.

We investigated PHA production, which represents a carbon storage mechanism, because defects in this function were previously reported for *ptsP* mutants in other organisms^36^. Our Δ*ptsP* mutant was indeed defective in PHA storage (Figure 6E), indicating that the PTS^Ntr^ impacts carbon storage in *P. aeruginosa*. Surprisingly, we observed an increase in PHA synthesis in a mutant of flagellar hook component *flgE*. Powering flagella requires a sustained investment of energy derived from the PMF, which itself is maintained by respiration and ultimately carbon catabolism. The increased PHA content of non-motile cells suggests that PHA storage and PMF maintenance are linked during growth arrest, and hints that carbon flux between these anabolic and catabolic processes could represent an important buffer *P. aeruginosa* (Figure 6F). Interestingly, studies in *E. coli* have suggested direct links between the PTS^Ntr^, flagellar motility and resource storage^47,48^. The PTS^Ntr^ system must also impact other aspects of cellular metabolism besides PHA storage, because mutations in *ptsP* and *ptsO* have strong fitness defects in multiple starvation conditions^20^ (Figure 5), while mutations in the PHA biosynthesis genes do not have strong effects under the starvation conditions we investigated. More work will be needed to gain a deeper understanding of how the PTS^Ntr^ system influences resource distribution in *P. aeruginosa*, including carbon storage via PHA.

Flagellar biosynthesis and use appear to play prominent roles in the expenditure of cellular resources during starvation in *P. aeruginosa*. Starved cells were observed swimming (Figure 1D), and our proteomics data suggest that flagellar synthesis continues during protracted starvation. An ongoing need for flagella is consistent with the recent report that *P. aeruginosa* continues swimming over long periods of carbon starvation, a tendency shared with a range of marine Gammaproteobacteria but not *E. coli*^26^. However, the preservation of flagellar motility also appears to have a cost, as its dysregulation through mutation of *fleN* imposed one of the strongest fitness defects we observed in our TnSeq data. In contrast, mutations of many of the flagellar structural components conferred fitness advantages, specifically across the transition between nitrogen and carbon starvation conditions. Presumably, some of these mutants still incurred costs of partial flagellar biosynthesis, so these fitness impacts may relate more to expenditures of the PMF by a fully functioning flagellum. Interestingly, two other genes that showed significant fitness effect across the starvation transitions we investigated were inhibitors of efflux pump expression, *mexR* and *nfxB* (Supplemental File 1). Mutations in these genes lead to overexpression of efflux pumps, which deplete the PMF^49^ and have previously been shown to cause loss of fitness under nutrient limitation^50^. Flagellar motility and efflux pumps can confer obvious fitness benefits that justify their costs to the PMF, in the presence of toxins that that must be removed from the cell or when better opportunities for colonization and growth exist at a distance. However, our data suggest that preserving PMF contributes substantially to fitness for *P. aeruginosa* in fluctuating growth arrested conditions (Figure 6F).

Much more work will be required to investigate the details of molecular mechanisms orchestrating the dynamic changes in activity and resource distribution we have observed in starved *P. aeruginosa*. Our experiments represent reductionist “end cases” with well-mixed planktonic cultures totally starved for nitrogen or carbon. This approach was used to probe the possibilities for ongoing biosynthesis under nutritional extremes, and to simplify the performance and interpretation of experiments. However, we propose that the same mechanisms are likely to contribute to buffering imbalances in resource availability in a range of non-steady-state growth conditions, which likely include complex natural habitats such as biofilms and human infection contexts. We hope that our genome-scale proteomic and genetic fitness data can serve as a resource to support future investigations of growth arrest regulation and physiology in *P. aeruginosa*.

## Materials and Methods

Please see supplemental materials and methods for additional details.

### Strains, Media, and Handling

MOPS minimal media lacking carbon and nitrogen sources (50mM MOPS, 40 mM NaCl, 4mM K2HPO4, 2.0 mM KCl, 1.0 mM MgSO4, 0.1 mM CaCl2,, 7.5 μM FeCl2·4H2O, 0.8 μM CoCl2·6H2O, 0.5 μM MnCl2·4H2O, 0.5 μM ZnCl2, 0.2 μM Na2MoO4·2H2O, 0.1 μM NiCl2·6H2O, 0.1 μM H3BO3, 0.01 μM CuSO4·5H2O) was used as a foundation for all nitrogen and carbon starvation experiments. 45 mM sodium succinate or 30 mM ammonium chloride were added as carbon or nitrogen sources respectively, as appropriate. *P. aeruginosa* UCBPP-PA14 and derived mutant strains (Supplemental Methods – Strain List) were consistently grown to stationary phase (∼24 hours) in lysogeny broth (10g/L tryptone, 5g/L yeast extract, 10g/L NaCl) at 37°C shaking prior to their resuspension in starvation media. Starved cultures were shaken and maintained at 37°C in all experiments.

To transition between starvations, cells were pelleted and washed at least once with pre-heated MOPS medium lacking carbon and nitrogen before resuspension in the new medium. These transfers could result in the loss of up to 50-75% of cells from cultures during transitions (Supplemental Figure 2A, B).

### Microscopy and Flow Cytometry

All images and videos were acquired using a Teledyne 48 photometrics camera fitted to a Nikon Eclipse Ti2 microscope with a 60X phase contrast oil objective (Nikon - MRD31605) and Spectra light engine (Lumencor). All microscopy images were processed in ImageJ using the microbeJ plugin^51^. Microscopy-based experiments which quantified cell area parameters involved data from 60-300 cells per biological replicate. For analyzing larger populations of cells, flow cytometry on either an LSR Fortessa (Becton Dickinson) or Novocyte (Agilent) flow cytometer was used. Cytometry data was imported and analyzed in FlowJo V10.8.1 (Becton Dickinson). Where required, fixed samples were stained with 1 μg/mL BODIPY (ThermoFisher) for 60 minutes at 37 C° in the dark and/or 10 μg/mL DAPI.

### Fluorescence in Situ Hybridisation

Cellular ribosome content was assessed using fluorescence in situ hybridisation as previously described,^27^ with minor modifications. Briefly, Cy5-labeled EUB338 16S rRNA or control “scramble” probes (Integrated DNA Technologies) were hybridized for 3 hours at 46°C in hybridization buffer (900 mM NaCl, 20 mM Tris pH 7.6, 0.01% SDS, 20% HiDi formamide and 2 μM fluorescent probe) against cells that had been fixed in 4% paraformaldehyde, permeabilized with ice cold ethanol and washed three times in 0.9% NaCl. Samples were then washed twice in wash buffer (215 mM NaCl, 20 mM Tris pH 7.6, 5 mM EDTA) for 15 minutes at 48°C shaking in the dark and resuspended in 0.9% NaCl for use in flow cytometry and microscopy. Scramble probe intensity was subtracted from EUB338 probe intensity during analysis.

### Fluorescence-Conjugated BONCAT Labelling

For whole cell BONCAT labelling, aliquots of culture were amended with azidohomoalanine (AHA, Tocris Bioscience), with labelling times and AHA concentrations indicated in figure legends. An AHA-free control was used for each condition and time point to determine background signal, with the average intensity of this control subtracted from labelled samples in analyses. Following labelling, cultures were pelleted using a benchtop centrifuge and washed three times in 0.9% NaCl to remove residual azidohomoalanine. 0.9% NaCl was used to dissolve all subsequent reagents and used to wash cells after each step. Cells were fixed with 4% paraformaldehyde at room temperature for 30 min, permeabilized in 70% ice cold ethanol, incubated with 100 mM iodoacetamide at 46°C for 30 minutes in the dark to alkylate free cysteines, incubated with 25 μM DBCO-AlexFluor-488 (Jena Bioscience) at room temperature for 30 minutes in the dark and were finally washed three times in 0.9% NaCl to remove residual dye for use in microscopy and flow cytometry measurements.

### Proteomics

For both proteomics experiments, cultures of UCBPP-PA14 were grown for 24 hours at 37°C shaking in LB before being normalised to an optical density of 0.2 in 100 - 300 mL nitrogen starvation MOPS minimal media and incubated/ sampled at time points as indicated (Figure 3).

### Total proteomics experiments

5 ml aliquots were flash frozen with liquid nitrogen and stored at -70 C°. Pellets were lysed in 2% SDS, 100 mM Tris pH 8.0, 0.2 μL/mL benzonase with cOmplete EDTA-free protease inhibitor (Sigma Aldrich), clarified by spinning at high speed in a benchtop centrifuge (13,000 g for 10 minutes) and normalised to a protein concentration of 150 μg/mL using lysis buffer. Trypsin digest, filter-aided sample preparation (FASP) and suspension trapping (sTRAP) were performed by the University of Dundee Fingerprints Proteomics facility. Peptides were loaded onto an Orbitrap Q-exactive plus LC-MS/MS instrument (ThermoScientific) with a 180-minute data-independent acquisition analysed via Spectronaut software (Biognosys) searching against the UCBPP-PA14 proteome (Uniprot reference UP000000653).

### BONCAT proteomics experiments

100 mL aliquots were removed from cultures and had AHA added to a final concentration of 500 μM for the time indicated in Figure 3. Following labelling, 200 µg/ml chloramphenicol was added to cultures to terminate protein synthesis before pelleting, washing in 0.9% NaCl and flash freezing. Cells were lysed as for total proteomics with the addition of 100 mM iodoacetamide and lysates normalized to equivalent protein concentrations. Urea buffer (8M Urea, 150 mM NaCl, 1X protease inhibitor) and 25 μL of DBCO agarose beads (Discovery Dyes) were added to normalised samples and rotated for four hours at room temperature. Beads were collected by centrifugation at 1000 rpm, washed with SDS wash buffer (0.8% SDS, 150 mM NaCl, 100 mM Tris pH 8.0) and resuspended in SDS wash buffer with 5 mM dithiothreitol at room temperature for 30. Beads were then incubated in SDS wash buffer with 100 mM iodoacetamide at 50°C in the dark for 45 minutes. Beads were washed in SDS wash buffer and transferred to Poly-Prep chromatography columns (BIO-RAD). Columns were washed 8 times with 5 mL SDS wash buffer, 8 times with 5 mL urea buffer with 100 mM Tris pH 8.0, and 8 times with 5 mL 20% acetonitrile. Finally, beads were resuspended in 10% acetonitrile with 50 mM ammonium bicarbonate. Peptides were recovered from the beads by tryptic digest, desalted, and analysed by LC-MS/MS as for total proteomes.

### Tn-Seq

Transposon insertion sequencing experiments were performed on two separate occasions, each using three biological replicates exposed to identical conditions and sampling time-courses (Figure 4.A). For both experiments, aliquots of the UCBPP-PA14 transposon mutant library (Supplemental Materials and Methods) were thawed and 50 μL was used to inoculate 50 mL of LB. Stationary phase samples (∼2mL) were taken after 24 hours growth, pelleted and frozen. The remainder of each LB culture was then pelleted, washed and resuspended to a final OD of 0.2 in 40 mL of either carbon starvation or nitrogen starvation MOPS minimal media. 0.5 OD units were collected after 48 hours incubation, washed, and resuspended in 5 mL LB for outgrowth. Following 2 hours of outgrowth, with cultures reaching an OD of ∼ 0.6, cultures were pelleted and frozen. The remaining cultures were split in half, pelleted, washed thoroughly in MOPS minimal media lacking both carbon and nitrogen sources, and resuspended in 10 mL of either nitrogen starvation or carbon starvation MOPS minimal media, achieving an OD of ∼0.05. Transitioned cultures were incubated for 24 hours at 37 C° shaking before sampling. The entire remaining cultures were pelleted and resuspended in 5 mL of LB for outgrowth. Following 2 hours of outgrowth, with cultures reaching an OD of ∼ 0.6, cultures were pelleted and frozen. Sequencing libraries were prepared as previously described^20^. Samples were then normalized, pooled, and sequenced on the Illumina NextSeq2000 instrument with a P1 reagent kit. Approximately 2.2 - 4.6 million single-end 100 bp reads were obtained per sample. Raw FASTQ files were processed for analysis using the Tn-Seq Pre-Processor (TPP) tool from the Transit package^52^, mapping reads to the UCBPP-PA14 genome (NC_008463.1). Read counts per gene were summarized using the FeatureCounts algorithm of the subread software package^53^. Both independent experiments were combined, so that each sampling timepoint is represented by at least five biological replicates (Supplemental File 1). The Voom/Limma differential expression method implemented on the Degust server was used to assess statistical significance of differences in read counts^54^.

## Acknowledgments

We are grateful for the *ΔptsP* and *ΔftsH* strains gifted to us by Dianne Newman. We appreciate assistance from the University of Dundee Flow Cytometry and Cell Sorting Facilty, Imaging Facility, and Fingerprints Proteomics Facility as well as the sequencing facility at the James Hutton Institute. We would like to thank Laurent Delavaine, Daniel Neill, Lisa Racki, and Dianne Newman for helpful feedback and discussions. Funding for this work was provided by the Wellcome Trust Institutional Strategic Support Fund to MB [204816/Z/16/Z], a Wellcome Trust PhD Studentship to FDM [218520/Z/19/Z]; the UK Academy of Medical Sciences (Springboard Award [SBF005/1096] to MB); and the UK Research and Innovation Medical Research Council (Future Leaders Fellowship [MR/T041811/1] and [MR/Z000378/1] to MB).

## Supplemental Information

**Supplemental Figure S1.**
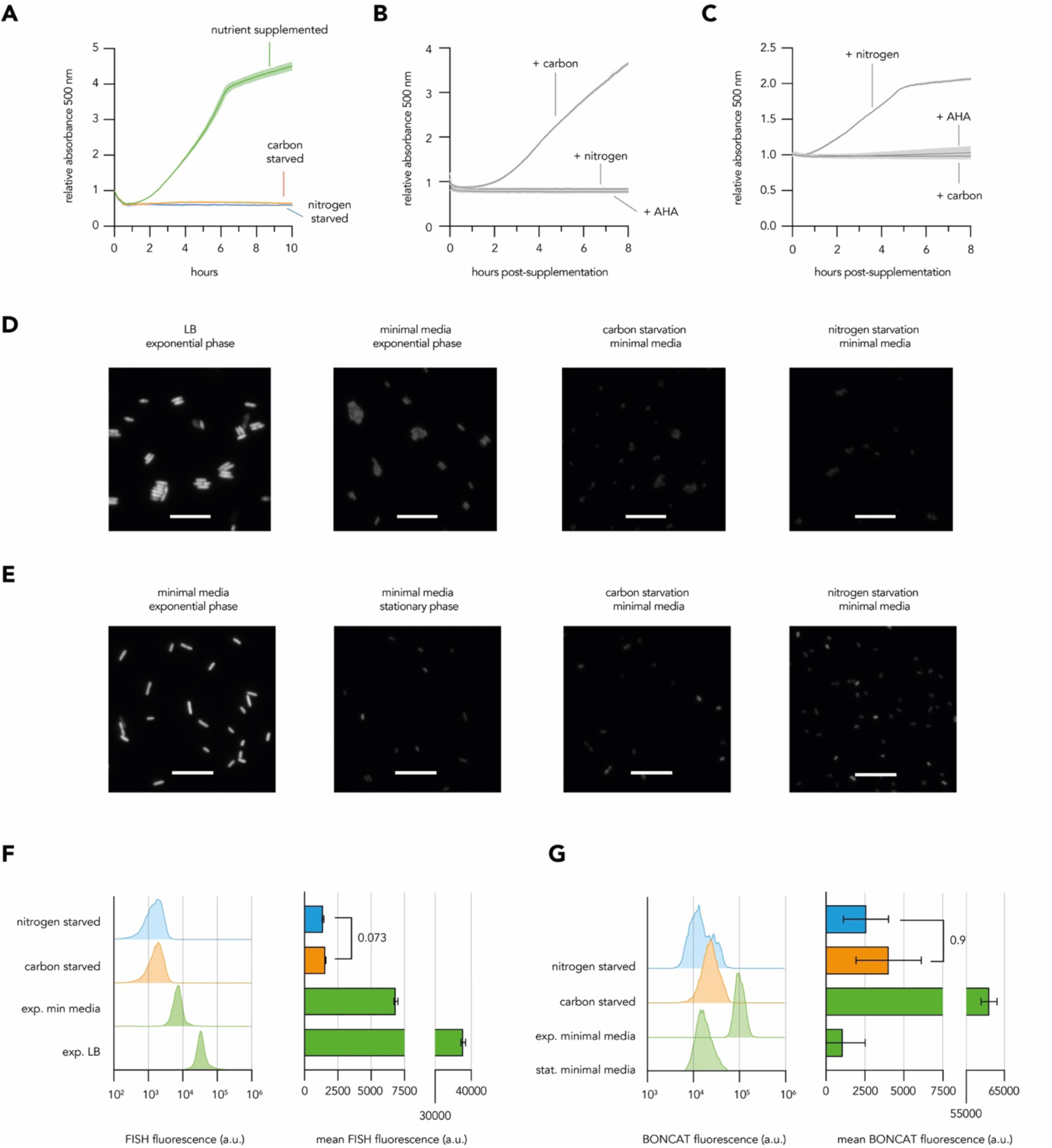
Line graphs (**A**) show the response of *P. aeruginosa* cultures to resuspension in nutrient-supplemented minimal media; carbon-starvation minimal media and nitrogen starvation minimal media. Resuscitation line graphs show the response of 48-hour carbon starved (**B**) and 48-hour nitrogen starved (**C**) *P. aeruginosa* cultures to supplementation with a carbon source (45 mM sodium succinate), a nitrogen source (15 mM ammonium chloride) and the amino acid analogue azido-homo-alanine (200 µM AHA). Representative fluorescence microscopy images show *P. aeruginosa* cultures which were fixed during growth-permitting conditions or following 50 hours of carbon or nitrogen starvation conditions and which were FISH-stained using a ribosome-directed probe (**D**) or BONCAT-labelled for 30 minutes with 500 µM AHA (**E**). In microscopy images, scale bar represents ten-microns. Histograms and bar graphs show the density of cellular fluorescence values and the mean cellular fluorescence values from *P. aeruginosa* cultures which were fixed during growth-permitting or starvation conditions and which were FISH-stained using a ribosome-directed probe (**F**) or BONCAT-labelled for 30 minutes with 500 µM AHA (**G**). All data represent the mean value calculated from at least three biological replicates with standard deviation displayed as error bars. Histograms are representative of at least three biological replicates. T-tests were used to calculate the p-values denoted above select comparisons, with only values close to or passing significance thresholds (p < 0.05) being shown. a.u. = arbitrary units.

**Supplemental Figure S2.**
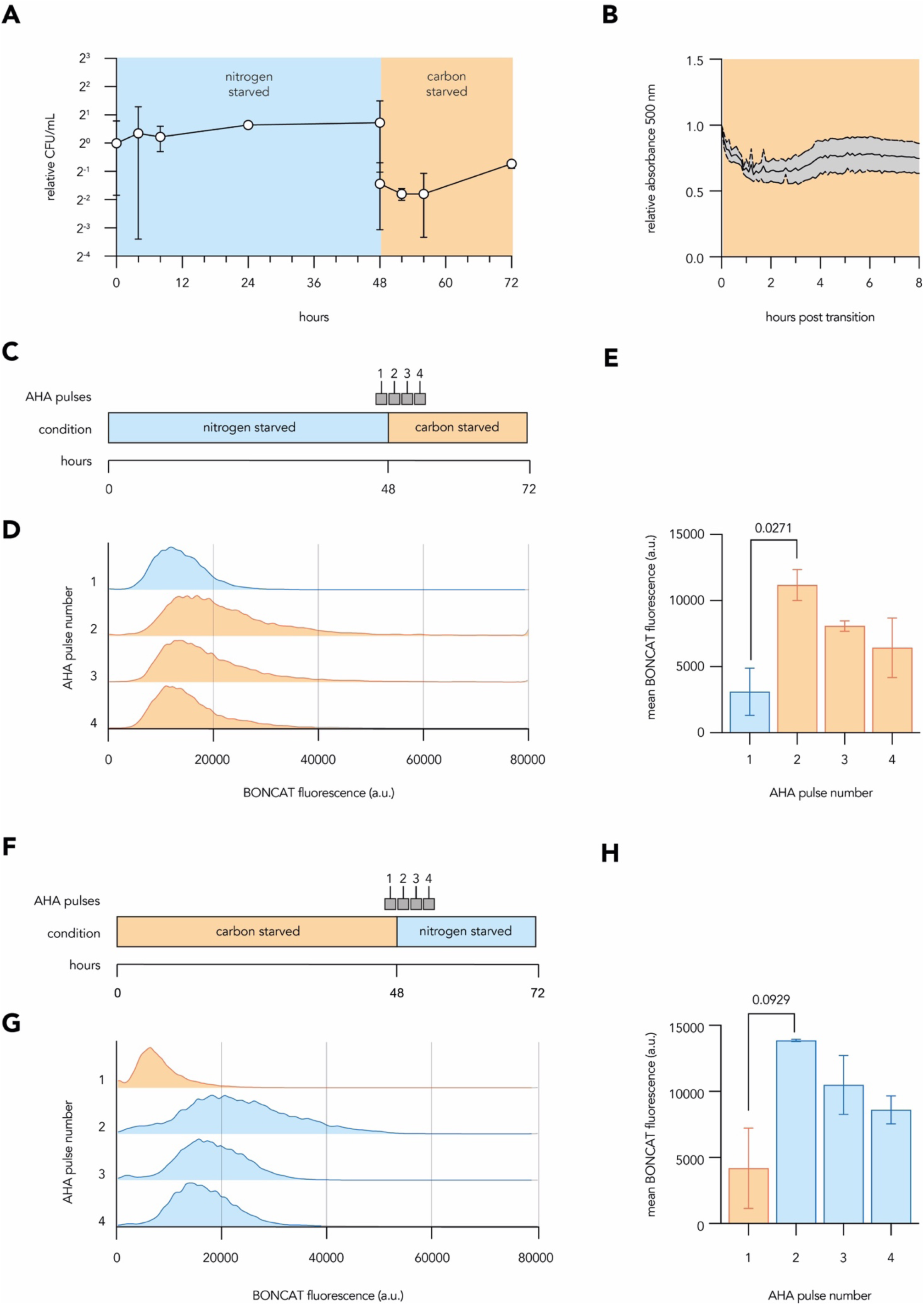
Line graphs show the trends in colony-forming units (**A**) and absorbance 500nm (**B**) during the rapid transition of *P. aeruginosa* cultures between nitrogen and carbon starvation minimal media. Schematics show the time-course and accompanying two-hour 500 µM AHA pulses applied to *P. aeruginosa* cultures during the BONCAT labelling of a transition from nitrogen starvation to carbon starvation minimal media (**C**) and the transition from carbon starvation to nitrogen starvation minimal media (**F**). Histograms (**D** and **G**) show the density of cellular fluorescence returned from each two-hour AHA pulse applied to transitioning cultures. Bar graphs (**E** and **H**) show the extracted mean fluorescence values from each BONCAT labelling pulse applied to transitioning cultures. All data represent the mean value calculated from at least three biological replicates with standard deviation displayed as error bars. Histograms are representative of at least three biological replicates. T-tests were used to calculate the p-values denoted above select comparisons, with only values close to or passing significance thresholds (p < 0.05) being shown. a.u. = arbitrary units.

**Supplemental Figure S3.**
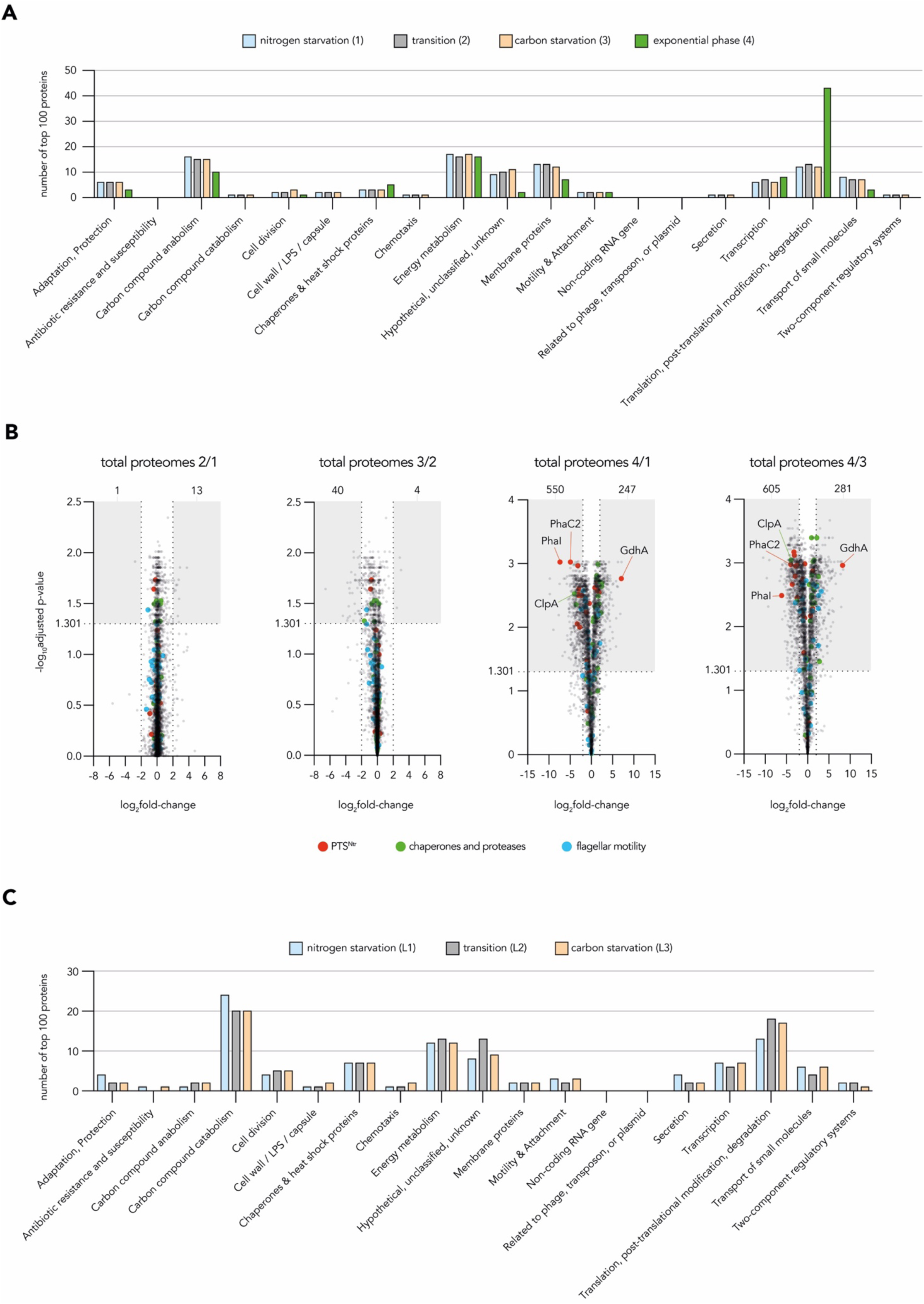
Bar graphs (**A**) show the occupancy of modified PseudoCAP groups by the top hundred most abundant proteins retrieved from the sampling timepoints of label-free proteomic experiments. Volcano plots (**B**) show the fold-change in abundance of proteins when comparing the total proteomes retrieved from different sampling periods of the label-free proteomic experiment. In each volcano plot, shaded boxes highlight the nascent proteins which passed fold-change (log_2_foldchange > 2 or < -2) and adjusted p-value (log_10_adjusted-p-value > 1.301) cut-offs to be considered as robust changes in abundance between compared proteomes, with the number of such proteins denoted above each shaded box. The larger coloured dots highlight individual proteins belonging to key physiological functions. Bar graphs (**C**) show the occupancy of modified PseudoCAP groups by the top hundred most abundant proteins retrieved from the labelling periods of BONCAT proteomic experiments outlined in Fig. 3D. All analyses used mean abundances calculated from four biological replicates.

**Supplemental Figure S4.**
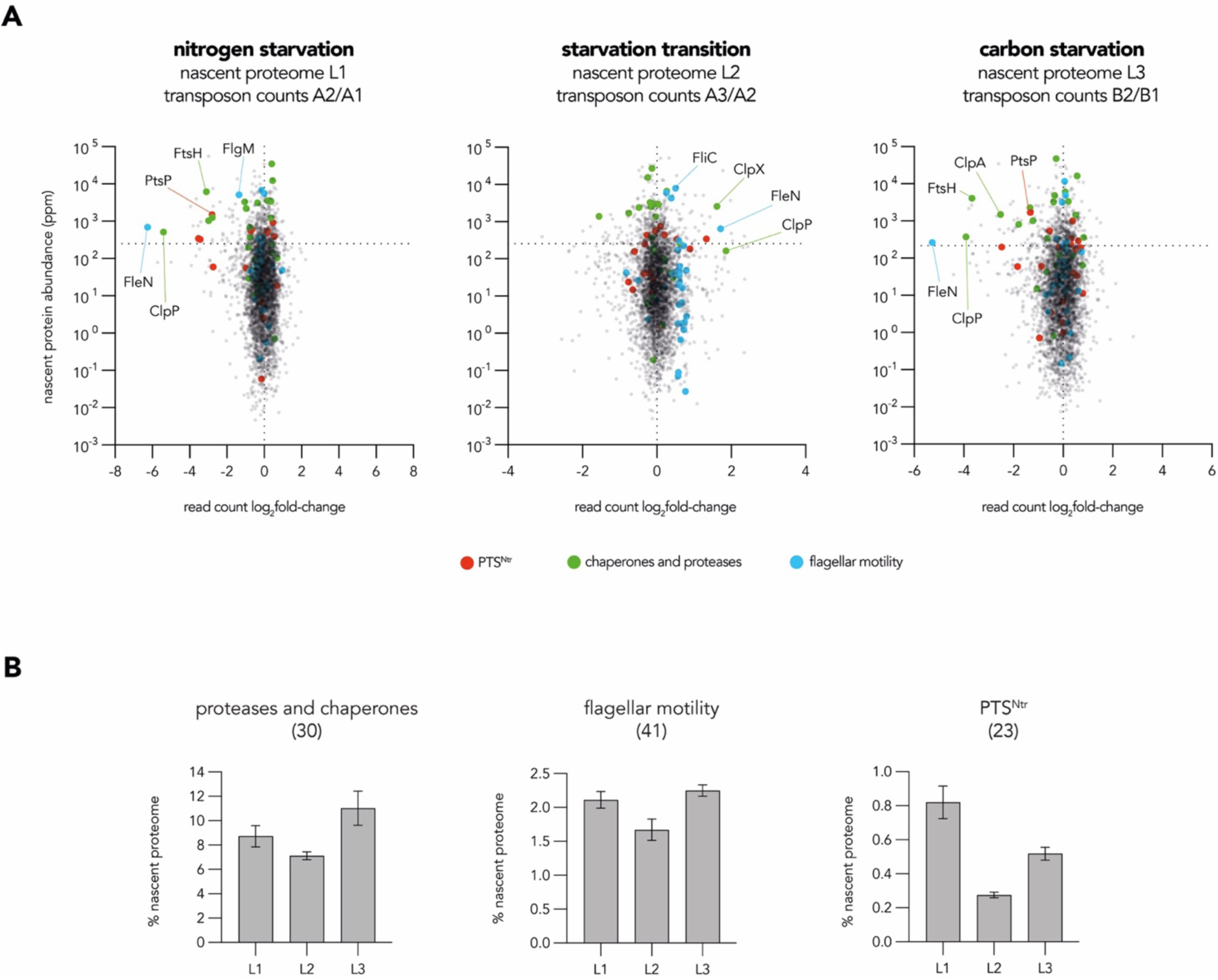
Scatter plots (**A**) combine BONCAT proteomic (outlined in Fig. 3D) and transposon-insertion sequencing data (outlined in Fig. 4A) and show the normalised protein abundance and transposon read count of individual proteins/genes detected in both datasets under comparable conditions. In each plot, a dotted line crosses the y-axis at values corresponding to the mean abundance of all normalised proteins within the refined proteomic dataset, and dotted line crosses the x-axis at x=0. The larger coloured dots highlight individual proteins belonging to key physiological features. Bar graphs (**B**) show the percentage occupancy of nascent proteomes by nascent proteins belonging to three key physiological features at each labelling period used in BONCAT proteomic experiments. The number of genes each discrete cohort is composed of are noted in brackets. All analyses used mean abundances calculated from four biological replicates of BONCAT proteomic data and mean read counts calculated from at least five biological replicates of TnSeq data. Bar graphs used the mean value calculated from four biological replicates, with standard deviation displayed as error bars.

**Supplemental Figure S5.**
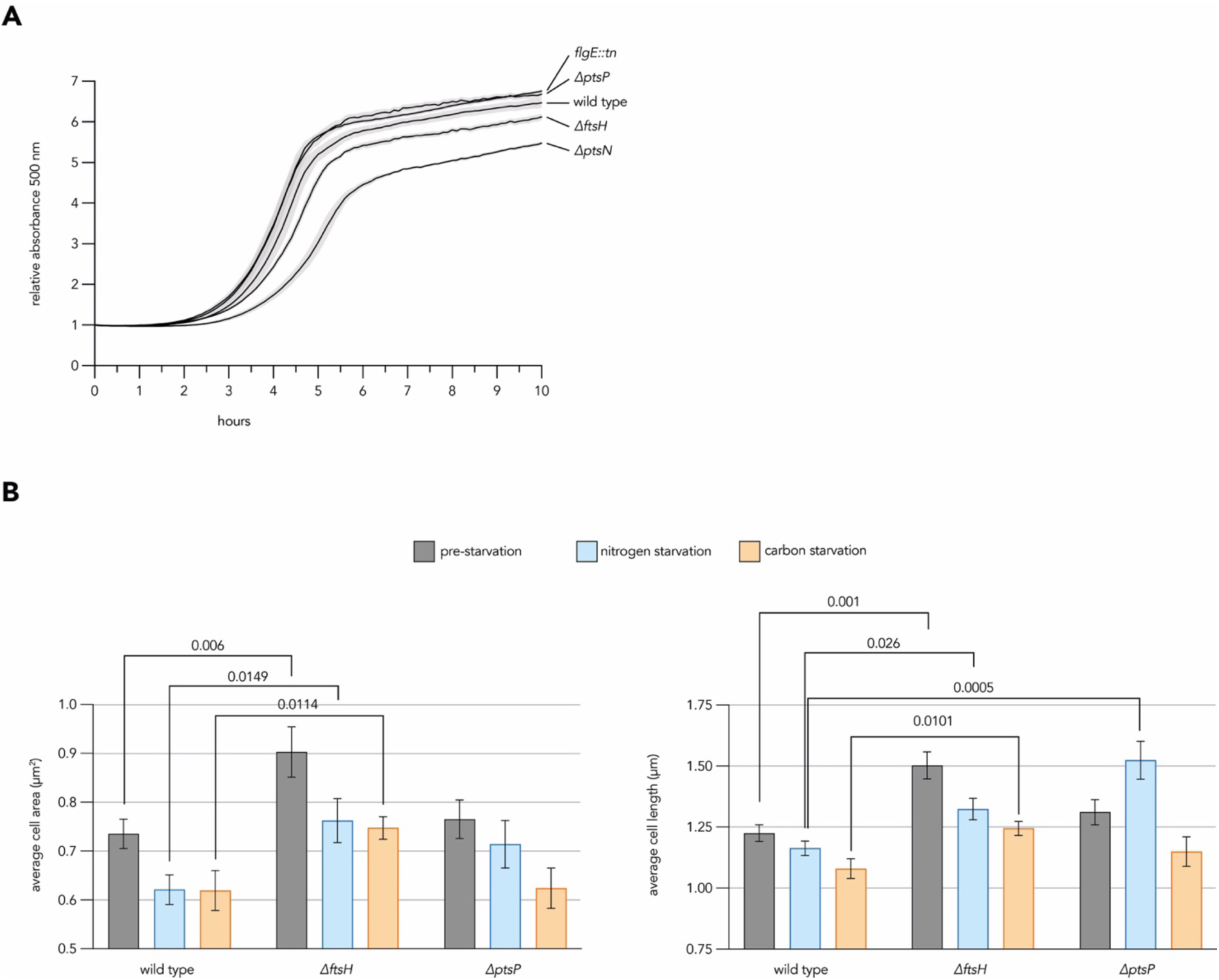
Line graph (**A**) shows the growth of mutant *P. aeruginosa* strains in LB media from a starting OD 0.01 as measured by absorbance at 500nm. Bar graphs (**B**) show the average cell area and cell length of fixed *P. aeruginosa* cells of different strains before resuspension in and following 48-hour incubation in nitrogen or carbon starvation media. All data represent the mean value calculated from at least three biological replicates with standard deviation displayed as error bars. T-tests were used to calculate the p-values denoted above select comparisons, with only values close to or passing significance thresholds (p < 0.05) being shown. a.u. = arbitrary units.

## Supplemental Materials and Methods

### Strain list

Genotypes and absence of background mutations were confirmed for all *P. aeruginosa* strains by whole-genome sequencing.

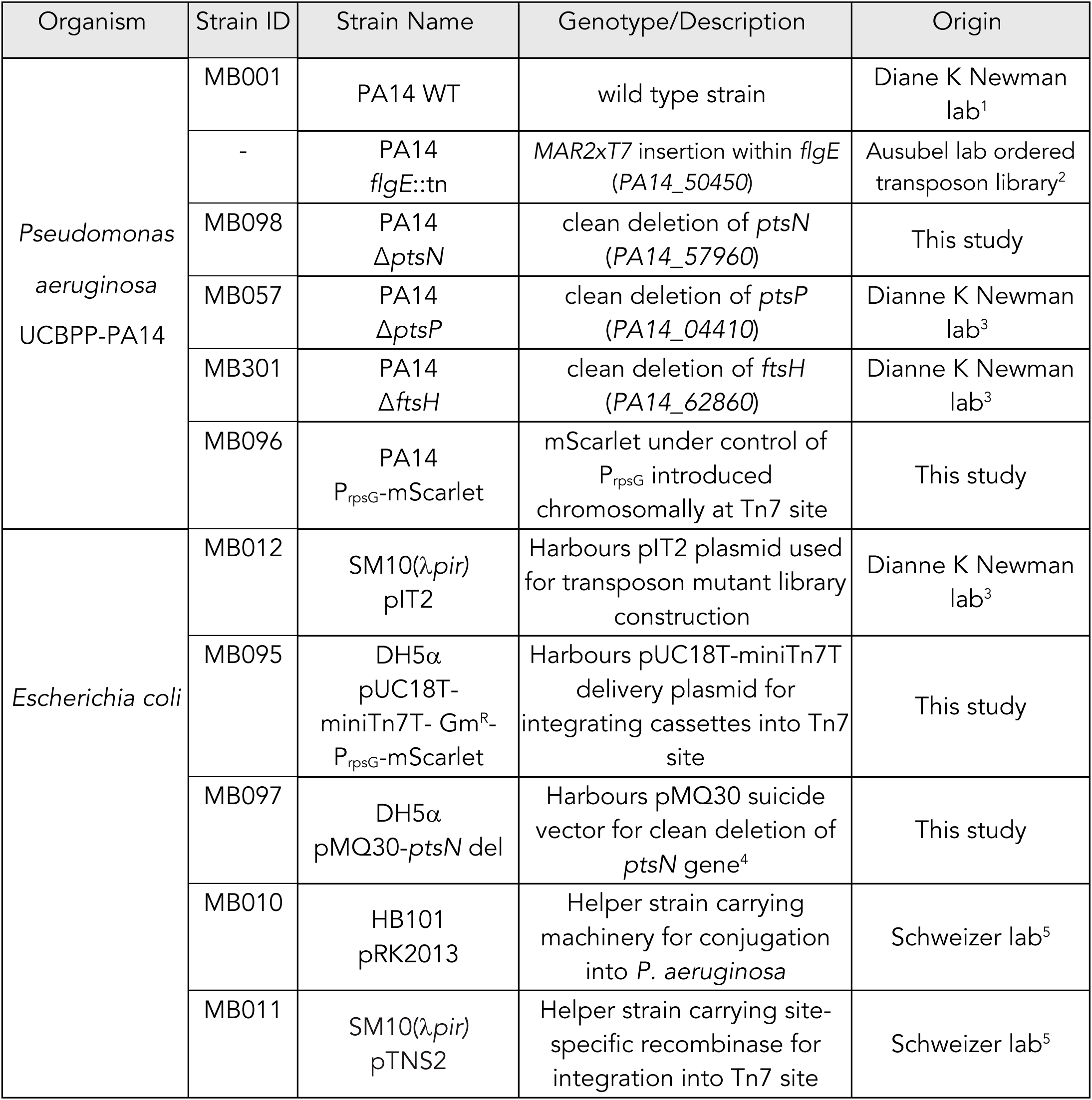

### Primer list

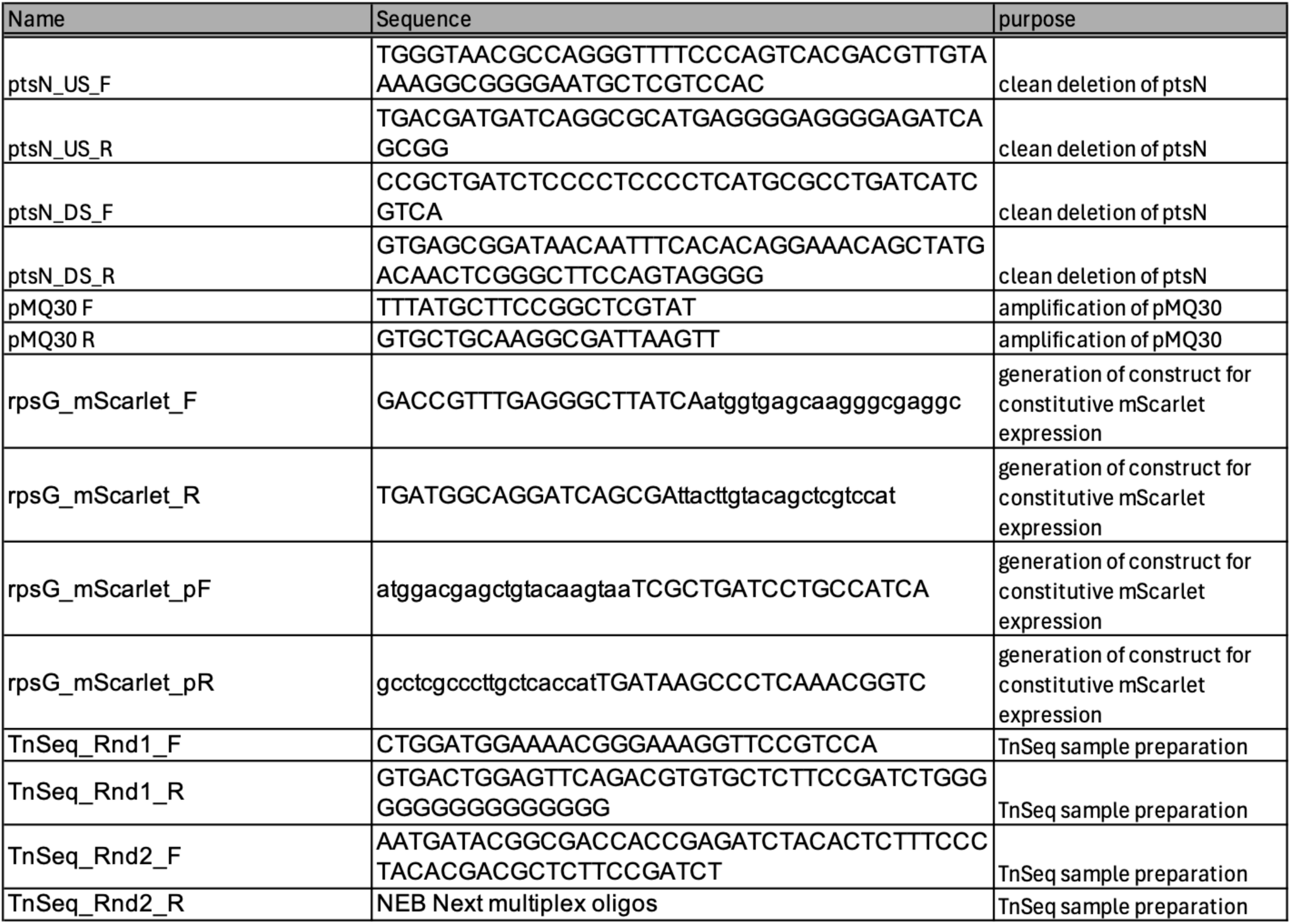

### Strain construction

An unmarked *ptsN* deletion strain (Δ*ptsN*) was produced by cloning two 600 bp sequences, encoding sequences upstream and downstream *ptsN* into the pMQ30 suicide vector^4^. These upstream and downstream regions were amplified from *P. aeruginosa* genomic DNA using primers containing additional sequences complementary to those of pMQ30. Gibson assembly was used to combine linearized pMQ30 plasmid (produced by PCR) with the amplified upstream and downstream fragments. Assembly product was used to transform competent *E. coli* DH5α cells, and successful transformants were selected on LB agar containing 100µg/mL gentamycin. The plasmid sequence of a successful transformant was confirmed by Sanger sequencing (MRC PPU DNA Sequencing and Services unit). Following confirmation, the pMQ30-*ptsN*del plasmid was introduced into *P. aeruginosa* UCBPP-PA14 by triparental conjugation. Successful exoconjugants were selected on VBMM medium (3 g/L trisodium citrate, 2 g/L citric acid, 10 g/L K2HPO4,3.5 g/LNaNH4PO4,1mM MgSO4, 100 μMCaCL2,pH 7) containing 100 μg/mL gentamicin as previously described^5^ and were then subjected to counterselection on LB plates lacking NaCl and containing 10% (wt/vol) sucrose. Colonies resulting from homologous re-combination to remove the wild-type copy of *ptsN* and retain the clean deletion were identified by PCR.

For the construction of the strain expressing mScarlet under the control of the *rpsG* promoter, the mScarlet-I encoding sequence was amplified from the pMRE-Tn7-145 plasmid^6^ and used to replace the sfGFP ecoding sequence in the plasmid from strain DKN1642^7^, by Gibson assembly. The mScarlet-ecoding cassette was integrated into the Tn7 site of the chromosome of WT PA14 using tetraparental conjugation, as described above but with the addition of the pTNS1-carrying helper strain^5^.

### Transposon-Insertion Sequencing – Library Preparation

The randomly inserting Tn5-based transposon delivery plasmid pIT2^8^ was conjugated into UCBPP-PA14 to produce a diverse transposon mutant library. UCBPP-PA14 and *E. coli* SM10(λ*pir)* carrying the transposon-bearing pIT2 plasmid were streaked onto LB agar plates and LB + 100 μg/mL carbenicillin agar plates respectively. Following overnight incubation at 37 C°, cells were scraped from each plate and resuspended in 1 mL of LB. Each suspension was then pelleted and washed once in LB before being resuspended in LB to an optical density of 50 and 100 OD units for PA14 and SM10(λ*pir)*/pIT2 respectively. Aliquots (100µL) of each dense culture were then mixed and 50 μL of this mixture was spotted onto LB agar plates and incubated at 37 C° for 2.5 hours. Spots were then scraped and resuspended in 16 mL of LB before 100 μL aliquots were spread onto 160 individual LB + 60 μg/mL tetracycline + 10 μg/mL chloramphenicol agar plates. Plates were incubated at 37 C° for 24 hours to select for transposon-integrated PA14. The roughly 180,000 resultant colonies were scraped and collected from the plates and made to an OD of ∼5 in LB. Glycerol was added to the pooled transposon mutant library which was then aliquoted and stored at - 70 C° until its application in subsequent experiments.

### Absorbance Readings

The absorbance of cultures at 500 nm was recorded using a Tecan Spark plate reader. At desired timepoints, aliquots of culture were placed in wells of clear-bottom 96-well plates and absorbance was recorded. In resuscitation assays, resuscitation was performed following prolonged incubation of cultures under starvation conditions. Samples (180μL) from aged cultures were aliquoted into wells of a flat-bottomed clear 96-well plate. Concentrated solutions of carbon, nitrogen or carbon + nitrogen were then added to wells to give a final volume of 200μL and a final concentration of 45 mM sodium succinate and/or 30 mM ammonium chloride. In experiments where azidohomoalanine was provided to cultures, the analogue was added to a final concentration of 500 μM. In resuscitation assays and growth curves, plates were sealed with a BreatheEasy gas-permeable sealing membrane (Diversified Biotech) and placed in a Tecan Spark plate reader pre-warmed to 37 C° and set to continuously shake. In all longitudinal experiments, absorbance readings were taken every 6 – 10 minutes.

### Agarose Pad Preparation for Microscopy

For each agarose pad, a 65 μL gene-frame (ThermoFisher) was attached to the surface of a standard 25 mm x 75 mm microscopy slide and used as an agarose holder. A 1% agarose solution was melted and ∼90 μL was added to a corner of the gene-frame. Immediately, a second slide was pressed on top of the gene-frame, forcing the agarose to completely fill the gene-frame. Gene-frame sandwiches were left to cool and solidify. The upper slide was then slid from the sandwich to reveal the solidified agarose. Sample droplets (0.5 μL) were spotted onto the agarose and allowed to dry before the gene-frame’s upper coating was removed and a cover slip was annealed to the exposed adhesive. Agarose pads were always imaged within a week of preparation and were kept at 4°C in the dark.

### Cellular Displacement Analyses of Bacterial Motility

Cultures of PA14 WT and PA14 *flgE*::tn were grown for 24 hours at 37 C° shaking in LB media before being washed and resuspended in nitrogen starvation or carbon starvation MOPS media at an OD of 0.2. Following 48 hours of incubation at 37°C, small samples were taken from cultures and delicately spotted onto the centre of a gene frame (ThermoFisher) attached to a glass slide. A coverslip was then adhered to the top of the gene frame, squashing the spot. Focusing just below the coverslip, samples were immediately filmed using the phase-contrast channel of a Nikon Eclipse Ti2 microscope. All films were 30 seconds in length and contained 101 frames (3 fps). The ImageJ plugin TrackMate was used to analyse phase-contrast films by identifying individual cells and tracking their displacement between frames of the film^9^. For each film, the data from cells which tracked for less than 50 frames or cells which exhibited extreme displacement indicative of faulty tracking were discarded. Cells were considered motile if they moved more than 4 microns during the 30 second films. The mean percentage motility of each strain was calculated from three biological replicate videos containing at least 40 cells.

### Transition Competition Assay

Cultures of PA14 P_rpsG_-mScralet and non-labelled strains of interest were grown for 24 hours at 37 C° shaking in LB media before being washed and resuspended in nitrogen starvation media at an OD of 0.2. Non-labelled competitor cultures, including a control of PA14 WT, were mixed in equal ratio with PA14 P_rpsG_-mScatlet and incubated at 37 C° shaking. Mixed cultures were incubated for 48 hours in nitrogen starvation MOPS minimal media before being transitioned to carbon starvation MOPS minimal media. Following another 48 hours of incubation, cultures were transitioned back to nitrogen starvation MOPS media. One more transition was applied, into carbon starvation MOPS minimal media, meaning that cultures had experienced a total of three starvation transitions. At the beginning and end of the time-course, small samples of these mixed cultures were taken for inspection via fluorescence microscopy. Fluorescence images of non-labelled and mScarlet-labelled cells were captured, and microbeJ^10^ was used to identify and quantify the number of each strain within the mixed cultures. The proportion of either strain within the mixture was calculated at the beginning and end of the time-course. The change in ratio of non-labelled mutant cells at the end of the time-course was then calculated relative to the ratio at the start of the time-course. For each mutant, this “competitive index” was then normalised to the value calculated in control competitions using non-labelled PA14 WT, which exhibited a small but consistent defect when competed against PA14 P_rpsG_-mScarlet. For each of three biological replicates used in competitions, quantification used images which contained between 90 – 400 cells.

### Microscopy and Image Analyses

All images and videos were acquired using a Nikon Eclipse Ti2 microscope fitted with a 60X phase contrast objective (Nikon - MRD31605); a TIPA slide adaptor (Oko Labs); a Nexus optical breadboard (Thorlabs); a camera (Teledyne 48 photometrics - SN A21b204004) and Spectra light engine (Lumencor - 80 10039). Fluorescence of interest was recorded with the following settings; BODIPY (excitation 470 nm LED, emission 510 nm filter cube); AlexaFluor-488 (excitation 470 nm LED, emission 510 nm filter cube); DAPI (excitation 395 nm LED, emission 435 nm filter cube); Cy5 (excitation 640 nm LED, emission 700 nm filter cube) and mScarlet (excitation 575 nm LED, emission 700 nm filter cube). Consistent phase-contrast settings were used across all relevant experiments, while fluorescence settings were calibrated to the brightest sample within each experiment.

All microscopy images were processed in ImageJ. Fluorescent channels were aligned to that of the phase using the ImageJ macro Align_Fluor_Channels, written by Norbert Vischer in 2017. The ImageJ plugin MicrobeJ was then used to identify and segregate individual cells and to quantify average cell shape parameters or average cellular fluorescence for BONCAT, FISH or BODIPY investigations^10^. Microscopy-based experiments which quantified cell area parameters involved data from 60-300 cells per biological replicate.

### Flow Cytometry Data Analyses

Fixed samples were analysed on either LSR Fortessa (Becton Dickinson) or Novocyte (Agilent) flow cytometers. Forward scatter pulse height (FSC-H) and side scatter height (SSC-H) parameters with logarithmic amplification were used to distinguish fixed bacterial cells from dust and other particulates, as was DAPI staining. Pulse height measurements were used for all fluorescence parameters. BODIPY and AlexaFluor-488 fluorescence was detected using 488 nm excitation and emission was collected at 530/30 nm on both LSR Fortessa and Novocyte cytometers. DAPI fluorescence was measured using 405 nm excitation and emission was collected at 450/50 nm and 445/45 nm on LSR Fortessa and Novocyte cytometers, respectively. Cy5 fluorescence was detected using 640 nm excitation and emission was collected at 670/14 and 675/30 on LSR Fortessa and Novocyte cytometers, respectively. Cytometry data was imported and analysed in FlowJo V10.8.1 (Becton Dickinson). Populations were first gated based on forward-scatter and side-scatter profiles to remove particulate before DNA-containing singlets were isolated by gating for DAPI profile. The resultant gated populations were used to produce fluorescence histograms in FlowJo and Adobe Illustrator, with the mean value of these histograms being directly extracted from FlowJo to provide and compare “average” fluorescent values of samples. Quantification of population average BODIPY, BONCAT and FISH fluorescence made use of between 7,000-40,000 cells per biological replicate.

### Modification of PseudoCAP and Choice of Key Features

For ease of data analyses and interpretation, established functional PseudoCAP^11^ were modified (Supplementary file 1). In cases where individual proteins had numerous PseudoCAP annotations, the first annotation was used. In cases where the first annotation was “hypothetical, unclassified, unknown” or “putative enzyme”, the second annotation was used. Several PseudoCAP categories were merged to simplify interpretation of data. The categories “amino acid biosynthesis and metabolism”; “biosynthesis of cofactors, prosthetic groups and carriers”; “central intermediary metabolism”; “fatty acid and phospholipid metabolism” and “nucleotide biogenesis and metabolism” were merged into “carbon compound anabolism”. The categories “cell division” and “DNA replication, recombination, modification and repair” were merged into “cell division”. The categories “protein secretion/export apparatus” and “secreted factors (toxins, enzymes, alginate) were merged into “secretion”. The categories “transcription, RNA processing and degradation” and “transcriptional regulators” were merged into “transcription”.

The key physiological functions we focused on (flagellar motility; proteases and chaperones and the phospho-transferase system) were defined and populated by selecting genes/proteins relevant to each (Supplemental file 1). For flagellar motility, members of the modified PseudoCAP category “motility & attachment” were largely used. Additional genes known to encode structural and regulatory flagellar components were also included. For proteases and chaperones, members of the modified PseudoCAP category “chaperones & heat shock proteins” were largely used. Additional genes known to encode proteases were also included. For the phospho-transferase system, genes encoding known components of the nitrogen-sensing and fructose-sensing phospho-transferase systems of *Pseudomonas aeruginosa* were included. Genes encoding metabolic enzymes whose products are predicted to influence these systems^12,13^ were also included. Genes relevant to the production of polyhydroxyalkanoate; an intracellular carbon storage device which is known to be influenced by phospho-transferase systems in *Pseudomonas putida*^14^ were also included.

### Total proteomics – Data Analysis

Data retrieved from total proteomic experiments were largely handled in Microsoft Excel and was assisted by described processing methodology^15^. For each sample, individual protein abundance was converted to parts per million to normalise for differences in processing and loading. The resulting values were considered reflective of the fraction of the total proteome represented by each protein, with non-detected proteins having an abundance set to 0. A principal components analysis of all samples was performed using the online tool ClustVis^16^. Refined data was uploaded to ClustVis, which returned numerous component values for each sample. For each sample, the first two components (PC1 and PC2) were extracted and plotted to confirm the clustering of replicates. In further analyses, the average abundance calculated from four biological replicates of each sampling timepoint was used. For abundance analyses, the top 100 most abundant proteins from each averaged sample timepoint were functionally classified into modified PseudoCAP (Supplemental file 1). Bar graphs of category occupancy were then produced for each sampling time point. For volcano plots, the fold-change for each detected protein was calculated using the average protein abundance for each individual sampling timepoint. Two tailed paired t-tests were used to calculate p-values for each comparison which were then adjusted using the Benjamini Hochberg method. Data were then transformed logarithmically, using log2(fold-change) and -log10(adjusted p-values), allowing to produce volcano plots (Supplemental file 1). Thresholds of substantial fold-change (log2(fold-change) greater than 2 or less than -2) and p-value (-log10(p-value) greater than 1.301) were applied to volcano plots to reveal proteins exhibiting robust differential abundance compared between sampling timepoints.

### Nascent Proteomics – Data Analyses

Data returned from nascent proteomic experiments were largely handled in Microsoft Excel and was assisted by described processing methodology^15^. For each sample, individual nascent protein abundance was converted to parts per million to normalise for differences in processing and loading, as well as differences in protein synthetic rates and labelling times. The resulting values can be considered to reflect the fraction of the newly synthesised proteome (produced during the labelling period) represented by each protein, with non-detected proteins having an abundance set to 0. A principal components analysis of all samples was performed using the online tool ClustVis^16^. Refined data was uploaded to ClustVis, which returned numerous component values for each sample. For each sample, the first two components (PC1 and PC2) were extracted and plotted to confirm the clustering of replicates. In further analyses, the average abundance calculated from four biological replicates of each sampling timepoint was used. For abundance analyses, the top 100 most abundant nascent proteins from each averaged labelling condition were categorised into modified PseudoCAP (Supplemental file 1). Bar graphs of category occupancy were then produced for each labelling condition. For volcano plots, the fold-changes for each protein were calculated using the average protein abundance for each individual labelling condition. Two tailed paired t-tests were used to calculate p-values for each comparison which were then adjusted using the Benjamini Hochberg method. Data were then transformed logarithmically, using log2(fold-change) and -log10(adjusted p-values), allowing to produce volcano plots (Supplemental file 1). Thresholds of substantial fold-change (log2(fold-change) greater than 2 or less than -2) and p-value (-log10(p-value) greater than 1.301) were applied to volcano plots to reveal proteins exhibiting robust differential expression compared between sampling timepoints. The average abundance of nascent proteins from three key physiological areas were extracted to produce heatmaps. For each nascent protein, the mean abundance retrieved from each individual labelling period was divided by the combined abundance across all three periods. These values of relative abundance were used in heat map production using Microsoft Excel.

### Transposon-Insertion Sequencing – Data Analyses

Pooled DNA libraries were prepared for sequencing on the Illumina NextSeq2000 instrument with a P1 reagent kit. Approximately 2.2 - 4.6 million single-end 100 bp reads were obtained per sample. Raw FASTQ files were processed for analysis using the Tn-Seq Pre-Processor (TPP) tool from the Transit package^17^, which finds and filters the transposon sequence from each read and maps the remaining genome sequence to a reference genome using BWA-MEM^18^. Approximately 2.5 - 5.0 million reads per sample were successfully mapped to the UCBPP-PA14 genome (NC_008463.1). Output.sam files were used in conjunction with the genome annotation to summarise counts per gene using the FeatureCounts algorithm of the subread software package^19^. Subsequent analyses used a combined dataset from two independent experiments, with each individual sampling timepoint represented by at least five biological replicates (Supplemental file 1). FeatureCounts data were uploaded Degust, an online tool for the exploration, analyses and visualisation of large sequencing datasets^20^. Analyses of differential read counts was then performed on select comparisons of interest using a Voom/Limma differential expression method (Supplemental file 1). In each select comparison these analyses returned fold-changes and FDR-adjusted p-values. Thresholds of substantial fold-change (log2(fold-change) > 0.585 or < -0.585) and false discovery rate (< 0.05) were applied to returned data to reveal genes exhibiting robust differential read counts between compared samples. These genes were defined as fitness-improving or fitness-diminishing determinants based on the directionality of the calculated fold change value. Genes whose transposon read count reduced in a defined comparison, suggesting that disruption of the gene reduced mutant fitness, were defined as fitness-improving. Genes whose transposon read count increased in a defined comparison, suggesting that disruption of the gene increased mutant fitness, were defined as fitness-diminishing. All substantial and significant fitness-improving and fitness-diminishing determinants returned from select comparisons were functionally categorised into modified PseudoCAP, allowing for bar graphs of category occupancy to be produced (Figure 4F).

### Combining Proteomic and Sequencing Datasets

Data retrieved from starvation and transition samples of label-free proteomic and BONCAT proteomic experiments were combined. Only proteins detected in both proteomic datasets were compared, with datasets being re-normalised following the removal of exclusive hits from either (Supplemental file 1). Any protein with an abundance of less than 1 ppm in either dataset was removed before scatter plots were produced. Proteins belonging to key groups of interest were then highlighted on these scatter plots. Data retrieved from starvation and transition samples/comparisons of BONCAT proteomic and transposon-insertion sequencing experiments were combined. Only proteins/genes detected in both nascent proteomic and transposon insertion sequencing datasets were compared, with the nascent proteomic dataset being re-normalised following the removal of exclusive hits (Supplemental file 1). Scatter plots incorporating both datasets were then produced, with proteins/genes belonging to key groups of interest were then highlighted on these scatter plots.

### Experimental Design and Statistical Rationale

Unless stated otherwise, experiments were performed in at least biological triplicate. In the context of frequently employed starvation experiments, biological replicates were defined as parental LB cultures derived from a distinct colony retrieved from fresh streak plates. For calculations of averages, the mean value of all biological replicates was always used, with the standard deviation of all biological replicates being used as error bars/ribbons. Statistical tests were often performed in GraphPad Prism V.10.1.0. T-tests and ANOVA tests were performed to infer significant distinctions between data in experiments. Trendlines were added to longitudinal BONCAT graphs using simple linear regression tools and non-linear fit tools for “binding saturation – one site total”. All figures and schematics were produced using a combination of Microsoft Excel, Microsoft PowerPoint, GraphPad Prism V.10.1.0, FlowJo V10.8.1 and Adobe Illustrator.

